# Highlighter: an optogenetic actuator for light-mediated, high resolution gene expression control in plants

**DOI:** 10.1101/2022.10.28.514161

**Authors:** Bo Larsen, Roberto Hofmann, Ines S Camacho, Richard W Clarke, J Clark Lagarias, Alex R Jones, Alexander M Jones

## Abstract

Optogenetic actuators have revolutionized the resolution at which we can assert control over biological processes in living systems. In plants, deployment of optogenetics is challenging due to the need for these light-responsive systems to maintain a single activation state in conventional horticultural environments with light-dark cycling. Furthermore, many available optogenetic actuators are based on plant photoreceptors that might crosstalk with endogenous signaling processes, while others depend on exogenously supplied cofactors. To overcome such challenges, we have developed Highlighter; a synthetic, light-gated gene expression system tailored for *in planta* function. Highlighter is based on the photoswitchable CcaS-CcaR system from cyanobacteria and is repurposed for plants as a fully genetically encoded system, engineered to photoswitch with the endogenous plant chromophore, phytochromobilin. We deployed Highlighter in transiently transformed *Nicotiana benthamiana* for optogenetic control of fluorescent protein expression and innate immune responses. Using light to guide differential fluorescent protein expression in nuclei of neighboring cells, we demonstrate unprecedented spatiotemporal control of target gene expression. We furthermore regulate activation of plant immunity by modulating the spectral composition of white light, demonstrating optogenetic control of a biological process in horticultural light environments. Highlighter is a step forward for optogenetics in plants and a technology for high-resolution gene induction that will advance fundamental plant biology and provide new opportunities for crop improvement.

## Introduction

Recent development of innovative and enabling high-resolution technologies has furthered the study of cellular processes, metabolic pathways, and regulatory systems. New measurements available to biologists, from single cell gene expression levels [1,2] to quantification of metabolites in tissues and in individual living cells [3,4,5], shines new light on the spatial and temporal relationships between quantified analyte and biological phenomena. However, to transcend the limits of correlative studies and establish causation, we must also be able to perturb biological systems with cellular-resolution.

Current tools for spatiotemporal perturbation, such as chemically inducible or tissue specific gene expression systems, can lack the desired resolution and may suffer from a series of additional limitations. For example, chemically inducible systems provide an element of temporal control, but typically depend on inducer molecules to diffuse into organs, tissues, and cells, limiting spatial and temporal resolution of application and removal. Further, they can be pharmacologically active, toxic, and expensive [6,7,8,9]. Correspondingly, cell-type or tissue-specific gene expression tools provide some degree of spatial control but are limited to previously characterized promoters and often lack specificity. However, optogenetic actuators, such as light inducible gene regulatory systems, could provide sought-after minimally-invasive, high-resolution control because light can be delivered with exquisite precision and with low toxicity.

One of the first reported synthetic light-controlled gene regulatory systems exploited a light-controlled protein-protein interaction between photoactive plant phytochromes and phytochrome interaction factors to drive reversible association of the split GAL4 transcription factor in *Saccharomyces cerevisiae* [10]. Following this breakthrough, the number of optogenetic actuator systems expanded rapidly from applications of light-controlled ion channels in neuroscience to numerous light-controlled biological processes in many cell types and even subcellular domains in living organisms [11]. Unfortunately, the implementation of optogenetic actuators in plants has proven challenging because plants require light-dark cycling for healthy growth and development. Most available optogenetic systems would be unable to maintain a single activation state under such conditions and thus applications are limited to those that can tolerate corresponding activation-inactivation cycles [11,12]. Furthermore, many optogenetic tools are based on light-responsive proteins from plants, such as PHYB, CRY2, PHOT&ZTL (LOV domains) and UVR8 [13], and may therefore crosstalk with endogenous light signaling pathways, potentially resulting in off-target modulation, or interference with the function of the synthetic optogenetic actuator itself. To minimize such problems, it is routine to orthogonalize system components (i.e. engineer system components not to interact with endogenous components) through mutation or truncation, as exemplified by the orthogonalized PhyB system [8]. Hence, ideal optogenetic actuators for plants will be (1) systems that specifically respond to artificial light stimuli, (2) assume a single activation state under standard plant growth conditions; i.e. light-dark cycling, (3) function as an optically controlled switch with distinct on- and off-states, (4) are orthogonal to plant signaling processes, and (5) do not require an exogenously supplied chromophore (see below).

In recent years, major advances have been made towards deploying optogenetic actuators to modulate gene expression in plants. Initially, an infrared-controlled actuator (infrared laser-evoked gene operator – IR-LEGO) [14] was deployed in plants to control gene expression from heat shock promoters with high resolution. However, the use of heat shock could lead to off-target gene induction in the targeted cells. Subsequently, a red light-controlled actuator, based on the N-terminal domains of PhyB and PIF6 from *Arabidopsis thaliana*, was demonstrated to achieve a high dynamic range of gene expression induction in *Nicotiana tabacum*- and *Physcomitrella patens*-derived protoplasts in response to 660 nm red light. However, 740 nm far-red light supplementation was needed to repress system activity under white light growth - conditions that affect endogenous phytochrome activity [8]. More recently, the bacterial green- and yellow-responsive CarH photoreceptor was developed as an optogenetic gene expression switch that responds to wavelengths of light that are minimally absorbed by plants [15]. This orthogonal optogenetic system was deployed in *Arabidopsis* protoplasts showing high induction and low background activity, but the photoactuator system is obligatorily dependent on the vitamin B_12_ derivative, adenosylcobalamin (AdoB12), an exogenously supplied proteolysis-sensitive chromophore. In a recent advance, the red-activated PhyB-PIF6 system was combined with an engineered blue-off module, based on the LOV-based transcription factor EL222 to generate a fully genetically encoded optogenetic gene expression system called PULSE. PULSE can be activated with red light when blue light is absent and remains off during light-dark cycling [16]. PULSE represents a major milestone for optogenetics in plants, as demonstrated by its deployment to reversible control induction of firefly luciferase expression in stably transformed *Arabidopsis*.

This work describes the design, engineering and validation of Highlighter; an optogenetic actuator tailored for regulating target gene expression levels in plants with cellular resolution. To make an optogenetic gene expression system that is orthogonal, fully genetically-encoded and independent of exogenously supplied chromophores, we chose to base the design of Highlighter on the CcaS-CcaR system, a green/red photoswitching transcription control system of cyanobacterial origin [17,18]. CcaS is a light-responsive histidine kinase that phosphorylates the response regulator CcaR, which then initiates transcription from a target promoter with cognate cis-regulatory elements (CREs) [17,18]. The CcaS-CcaR system was previously repurposed into synthetic optogenetic gene expression systems for prokaryotic hosts such as *Escherichia coli* [19,20,21,22,23,24], *Bacillus subtilis* [25] and cyanobacteria [26,27,28]. Target genes were placed under control of promoters with CcaR CREs and biosynthetic genes for the native chromophore of CcaS, phycocyanobilin (PCB), were exogenously expressed in hosts not naturally producing this chromophore. When repurposing this system for deployment in plants, we hypothesized that CcaS, having homology to plant phytochromes, might accept the endogenously produced phytochromobilin (PΦB) chromophore, which supports photoswitching in plant phytochromes [29]. It was expected that PΦB substitution for PCB in the green/red cyanobacteriochrome CcaS would generate a functional analog and hence circumvent the need for exogenously supplied chromophores. Moreover, the light environment used to sustain robust plant growth might be suitably adjusted to maintain the CcaS system in the same activity state in both the light and dark phases of diurnal growth. Light regimes artificially enriched in green light could then be used to activate this system with minimal perturbation of endogenous signaling processes, or photosynthesis itself, as these are less responsive to green light. By repurposing a system of prokaryotic origin we potentially also minimize crosstalk between the optogenetic actuator and endogenous plant signaling pathways.

We engineered Highlighter to function in eukaryotic cells and to efficiently photoswitch with PΦB by mutating the chromophore binding domain in CcaS, thus enabling use in higher plants that naturally synthesize PΦB. We found that target gene expression levels can be specifically repressed with blue light and blue-enriched white light, and activated with other light regimes, e.g. green-enriched white light. In *Nicotiana benthamiana* leaves transiently expressing Highlighter, we demonstrated robust optical control over fluorescent protein expression levels and induction of immune responses. We furthermore demonstrate the exquisite spatiotemporal control afforded by optogenetic actuators by using Highlighter to drive contrasting expression states in neighboring cells. Because target gene expression can be modulated by altering the spectral properties of white light, Highlighter’s behavior presents a solution for achieving minimally-invasive regulation of target gene expression levels under standard horticultural light regimes without the need to combine systems with opposing properties. Highlighter therefore provides new opportunities for optogenetic perturbation of biological processes with high spatiotemporal resolution in plants.

## Results

The primary challenge of developing Highlighter was to repurpose the CcaS-CcaR system for target gene control in plants, i.e. a eukaryotic host, with the endogenously produced PΦB chromophore. We envisioned that if PΦB supports photoswitching in CcaS, then efficient target gene control *in planta* could be achieved by targeting CcaS-CcaR to the plant nucleus with nuclear localization signals (NLS), codon-optimizing the system for plant expression, adding eukaryotic trans-activation domains (TADs) to CcaR and engineering a cognate synthetic promoter that incorporates recognition sequences for CcaR adjacent to a minimal plant promoter. Light-activation of the re-engineered CcaS-Highlighter (CcaS_HL_), would in principle, activate the optimized CcaR-Highlighter (CcaR_HL_), which would bind to the synthetic Highlighter promoter (P_HL_) and recruit the eukaryotic transcriptional machinery – resulting in target gene expression (Fig 1).

**Fig 1.**
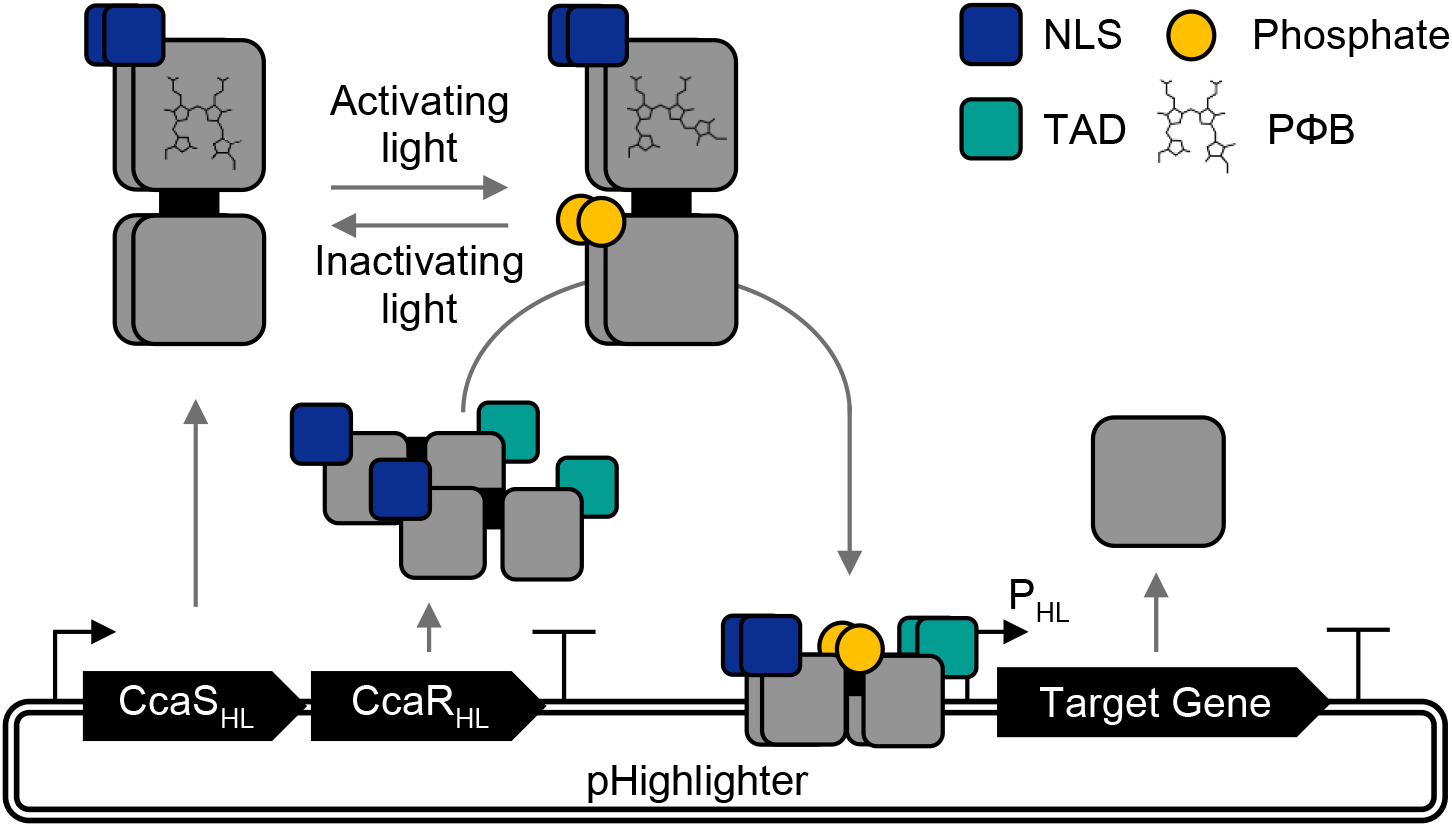
Schematic representation of the Highlighter system and function. Highlighter is the CcaS-CcaR system repurposed for *in planta* function. The repurposed CcaS, CcaR and synthetic promoter are denoted with subscript “HL”. Upon exposure to activating light conditions, CcaS_HL_ phosphorylates CcaR_HL_, which triggers enhanced binding to its cognate promoter, P_HL_, to induce expression of a target gene of interest. CcaS_HL_ and CcaR_HL_ are expressed as a single transcriptional unit from a promoter-terminator expression cassette through use of a F2A_30_ ribosomal skipping sequence. NLS, nuclear localization signal; TAD, transcription activation domain.

### Characterizing chromophore compatibility of the CcaS-CcaR system with P?B

We first set out to confirm if CcaS can photoswitch effectively with the endogenously produced plant chromophore PΦB. PΦB and the native PCB chromophore of CcaS are structurally similar heme-derived linear tetrapyrroles, which differ by exchange of the 18-ethyl group of PCB with an 18-vinyl group in PΦB [30]. Plant phytochromes PhyA and PhyB have previously been demonstrated to be able to photoswitch with both chromophores [31,32,33]. To test the chromophore dependency of CcaS we used a CcaS-CcaR system variant repurposed for *E. coli*, where CcaS and CcaR are expressed together with two cyanobacterial enzymes, HO1 (heme oxygenase) and PcyA (ferredoxin-dependent bilin reductase), to synthesize PCB from heme [34,19,20]. To report system activity, superfolder green fluorescent protein (sfGFP) is under the control of an optimized CcaR promoter, P_cpcG2-172_ [20]. Side-by-side comparison of system activity with PCB and PΦB was achieved by substituting *pcyA* with *mHY2*, which encodes a PΦB synthase from *Arabidopsis* lacking its native transit peptide [33].

Expressing the CcaS-CcaR reporter system in PCB-producing *E. coIi* cultures yielded a green/red switching transcription actuator system, as expected, which operationally can be activated by wavelengths in the visible spectrum shorter than 630 nm and be repressed by wavelengths greater than 630 nm (Fig 2A). By contrast, in PΦB-producing *E. coIi* cultures, sfGFP expression was not robustly light regulated, suggesting that CcaS poorly binds PΦB or does not photoswitch as well as the PCB adduct in *E. coli*.

**Fig 2.**
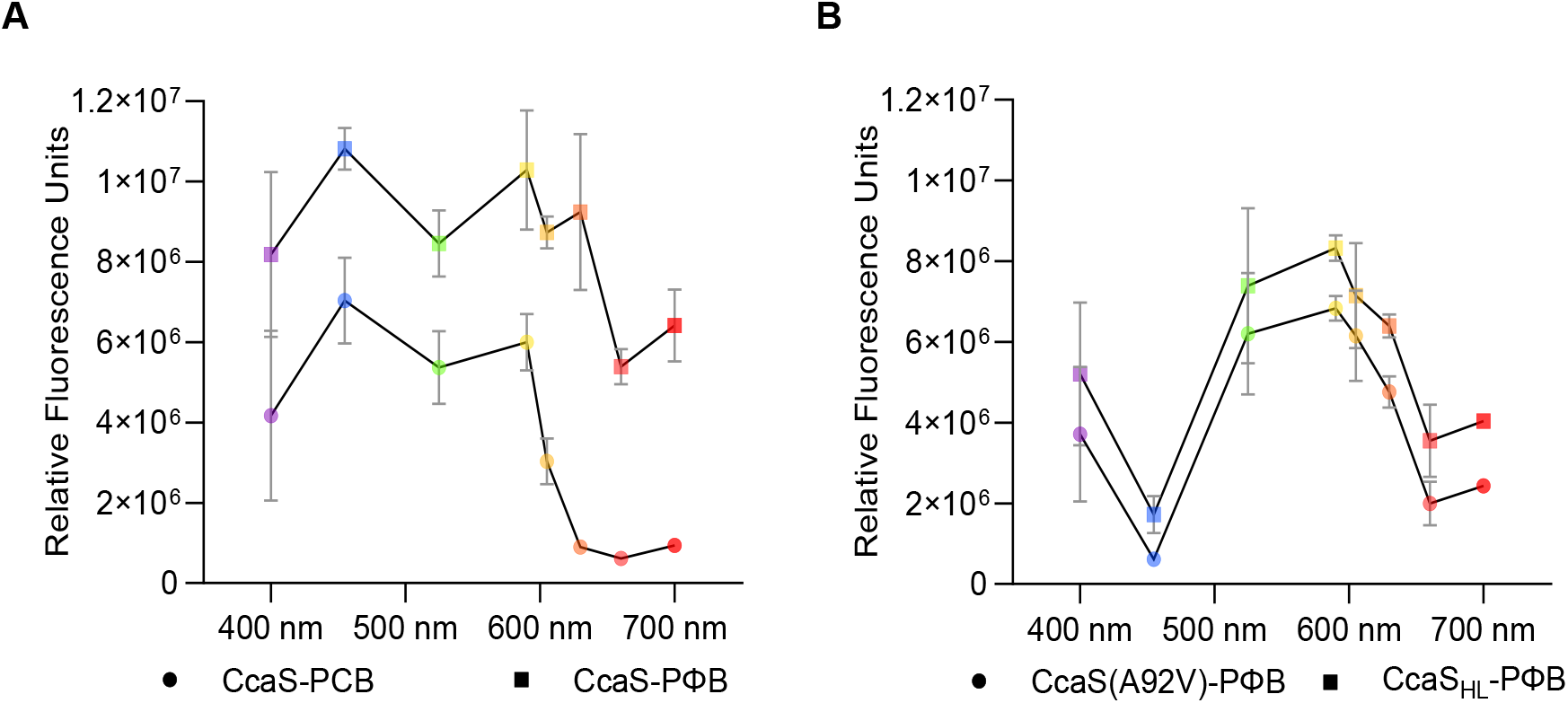
Chromatic response of CcaS-CcaR system variants in *E. coli*. (A) Chromophore-dependent photoswitching behavior of the CcaS-CcaR system in *E. coli* with PCB and PΦB with unmodified CcaS and CcaR proteins. System output in response to light stimuli was quantified via sfGFP fluorescence and presented as relative fluorescence units (RFU); RFU are defined as the mean estimated sfGFP fluorescence from cell cultures with OD600 nm = 0.2. Bacterial cultures were exposed to light stimuli (∼10 μmol m^−2^ s^−1^) generated using light emitting diodes (LEDs) with peak wavelength emissions around 400 nm, 455 nm, 525 nm, 590 nm, 605 nm, 630 nm, 660 nm and 695 nm. (B) System responses for CcaS(A92V) and CcaS_HL_ co-produced with PΦB. Symbols are colored according to light treatment and depict mean fluorescence from 3 replicate experiments, each comprising 3 biological replicates for each light treatment. S.E.M are presented for each light treatment.

### Improving photoswitching of CcaS with PΦB

To functionally tune CcaS for efficient photoswitching with PΦB, we selected residues for mutagenesis in and around the chromophore binding pocket of CcaS: specifically, conserved amino acid residues within the chromophore-binding cGMP phosphodiesterase/adenylyl cyclase/FhlA (GAF) domain of CcaS homologs that utilize PΦB. Sequence alignments for cyanobacteriochromes (NpR6012g4, TePixJ, FdRcaE, SyCcaS and SyCph1), plant phytochromes (AtPhyA and AtPhyB) and bacteriophytochromes (PsBphP and DrBphP) revealed candidate amino acids potentially associated with PΦB utilization (S1 Fig). The A92V mutation in the CcaS-GAF domain dramatically improved photoswitchable transcriptional regulation with PΦB in *E*.*coli* (Fig 2B). CcaS(A92V) with PΦB exhibited green/red regulation, similar to wild-type CcaS with PCB, but the former appears to require slightly longer wavelengths of light, i.e. red and far-red light from 660 nm, to switch off sfGFP expression. A more profound difference in spectral response was the blue-off behavior of CcaS(A92V) with PΦB, not seen for the unmodified CcaS with PCB. Blue light treatment with wavelengths around 455 nm efficiently lowered system output levels, as reported via the diminished sfGFP fluorescence.

### Modifying CcaS for function in plants

The next step towards repurposing the CcaS-CcaR system for efficient transcription control in plants was to target CcaS(A92V) and CcaR to the plant nucleus by fusing NLS domains to both proteins. For CcaS, this entailed removal of the N-terminal transmembrane domain (TMD) to release the photoreceptor from the cell membrane. We tested in *E. coli* whether replacing the TMD with an NLS produced a viable CcaS(A92V) before continuing to modify the system for deployment *in planta*. We found that CcaS(A92V) with the NLS substitution, hereafter CcaS_HL_, with PΦB had a slightly reduced dynamic range of response, compared to CcaS(A92V) with PΦB, but that the modification did not appear to further change the photoreceptor’s photoswitching properties (Fig 2B).

To investigate the response of CcaS_HL_ with PΦB to different illumination wavelengths, we heterologously expressed and purified a hexahistidine-tagged CcaS_HL_ holo-protein from *E. coli*. Spectroscopic data confirm that the recombinant PΦB adduct of CcaS_HL_ is reversibly photoswitched between red-absorbing (active) and green-absorbing (inactive) states (S2A, 2B, 2C and 2D Fig), similar to PCB adducts of the CcaS-GAF domain and truncated CcaS (CcaS without the transmembrane domain) expressed in *E. coli* or in *Synechocystis* sp. PCC 6803 [17]. By comparison with PCB adducts, photoconversion of the recombinant PΦB adduct of CcaS_HL_ between each state appears less complete. For simplicity we will continue to refer to them as the red- and green-absorbing states, but they can be more fully described as ‘red/green-absorbing’ and ‘predominantly green-absorbing’, respectively, an important distinction to some of the discussion below. An isosbestic point exists at ∼ 605 nm between these states, consistent with the transition between the active and inactive optogenetic behavior we observe in *E. coli* (Fig 2b). Blue light illumination has the same effect on the absorption spectrum as green light (S2c and S2f Figs); i.e., it converts the green-absorbing to the red-absorbing state. This suggests blue light should switch CcaS_HL_ into the active state, which appears to contradict the observed effect of blue light in *E. coli* (Fig 2b), where it switches off sfGFP expression.

Could this effect of blue light be explained by another compelling difference between the spectra of the PΦB adduct of CcaS_HL_ and the PCB adduct of CcaS GAF domain [17]? We observe additional, unexpected absorption peaks at around 475 nm and 445 nm, which resemble signal from oxidized flavin partially obscured by the UVA absorption peaks from PΦB. Together, these two peaks in the blue have the sort of defined vibrational structure observed when oxidised flavin is protein-bound (e.g., LOV2 from phototropin, [35]). Moreover, the magnitude of the UVA peak we observe for the green-absorbing state of the PΦB chromophore in CcaS_HL_ is greater relative to the green peak than one might expect [17], consistent with additional absorption in this region from oxidized flavin. It is therefore possible that oxidized flavin, e.g. flavin mononucleotide (FMN), is bound to the PAS domain of CcaS_HL_, similar to LOV domains. To investigate this further, we acquired the fluorescence emission spectrum (S3 Fig), exciting the red-absorbing state (from which blue light causes no further photoisomerization of the PΦB, S2f Fig) near the peak of the putative flavin absorption signal (445 nm). Because fluorescence signals typically come from the lowest-lying excited state, emission from flavin would not be obscured by transitions associated with higher-lying states of the PΦB. This is evident in S3 Fig, where a broad emission with peaks at 495 nm and 515 nm is, like the visible absorption signals, strongly reminiscent of the equivalent FMN spectrum from the phototropin LOV2 domain [35]. The peaks at 628 nm and 666 nm are likely from transitions associated with the residual broad green absorption in the red-absorbing state of the PΦB chromophore in the GAF domain of CcaS_HL_. Interestingly, the second PAS domain of CcaS exhibits high sequence similarity to LOV2 domains, for example 39.8% identity to the LOV2 domain of *Arabidopsis* PHOTOTROPIN 1. The PAS domain in CcaS has several of the key flavin-coordinating residues observed in flavin-binding LOV domains, e.g. CcaS G_433_KTPRVLQ_440_ motif, excepting the conserved cysteine that is involved in photochemical flavin-adduct generation in LOV domains. It is possible that the counterintuitive effect of blue light on CcaS_HL_ in *E. coli* (Fig 2b) is owing to a non-canonical effect on flavin bound to this cysteine-less, LOV-like domain, as previously shown for several other LOV receptors [36].

### Deploying Highlighter in transiently transformed *N. benthamiana* for optogenetic control of target gene expression

Having engineered CcaS_HL_ for nuclear targeting and to photoswitch with PΦB, we optimized the CcaS-CcaR system for plant deployment. First, to regulate target gene expression levels *in planta*, we needed to enable CcaR to effectively recruit the plant transcriptional machinery and initiate transcription, and second, we needed to create a plant compatible promoter recognized by CcaR. To achieve this, we converted CcaR into an eukaryotic transcription factor by adding a synthetic, C-terminal VP64 transcription activation domain [37] and for efficient activation by nuclear targeted CcaS_HL_ we included an N-terminal NLS domain (the resulting nlsCcaR:VP64 is hereafter referred to as CcaR_HL_). In parallel, we designed a synthetic cognate promoter for CcaR_HL_, named P_HL_, by placing three CcaR binding motifs 5’ to a 35S minimal promoter [38]. The binding motifs were spaced evenly around the DNA helix, offset relative to one another at approximately 120° angles, to maximize the chance of having at least one P_HL_-bound CcaR_HL_ being advantageously oriented to the 35S minimal promoter. Finally, considering that both CcaS and CcaR are of prokaryotic origin, we codon-optimized CcaS_HL_ and CcaR_HL_ for plant expression. Together, these repurposed components comprise the Highlighter system.

To demonstrate optogenetic control of target gene expression with Highlighter *in planta*, we constructed a series of vectors for deployment in transiently transformed *N. benthamiana*. Because heterologous gene expression levels can be variable in transient expression systems, we devised a ratiometric fluorescent reporter system for tracking system activity. To report target gene regulation we placed nlsedAFPt9 [39], a nuclear targeted yellow fluorescence protein (YFP) variant, under P_HL_ control, creating a YFP response module (P_HL_::nlsedAFPt9::T_nos_). For normalization of target gene expression levels to system gene expression levels we transcriptionally linked expression of the reporter protein nlsTagRFP, a nuclear targeted red fluorescence protein, with constitutive CcaS_HL_ and CcaR_HL_ expression. Transcriptional linkage of CcaS_HL_, nlsTagRFP and CcaR_HL_ from a single promoter-terminator cassette was achieved with two F2A_30_ ribosomal skipping sequences [40], effectively creating the Highlighter expression module (P_UBQ10_::CcaS_HL_:F2A_30_:nlsTagRFP:F2A_30_:CcaR_HL_::T_RBCS_). Together, these two modules comprise the Highlighter(YFP) system.

In *N. benthamiana* leaves infiltrated with *Agrobacterium tumefaciens* for transient transformation with Highlighter(YFP), we compared the ratio of nuclear YFP/RFP fluorescence under different light conditions. We observed markedly lower YFP/RFP ratios in samples kept in continuous blue light (λ ∼ 455 nm), indicating lower relative target gene expression levels, as compared to leaves subjected to green light (λ ∼ 525 nm) or red light (λ ∼ 660 nm) (Fig 3A and 3B). Target gene expression levels were verified to correlate with the observed fluorescence ratios by qRT-PCR (Fig 3C).

**Fig 3.**
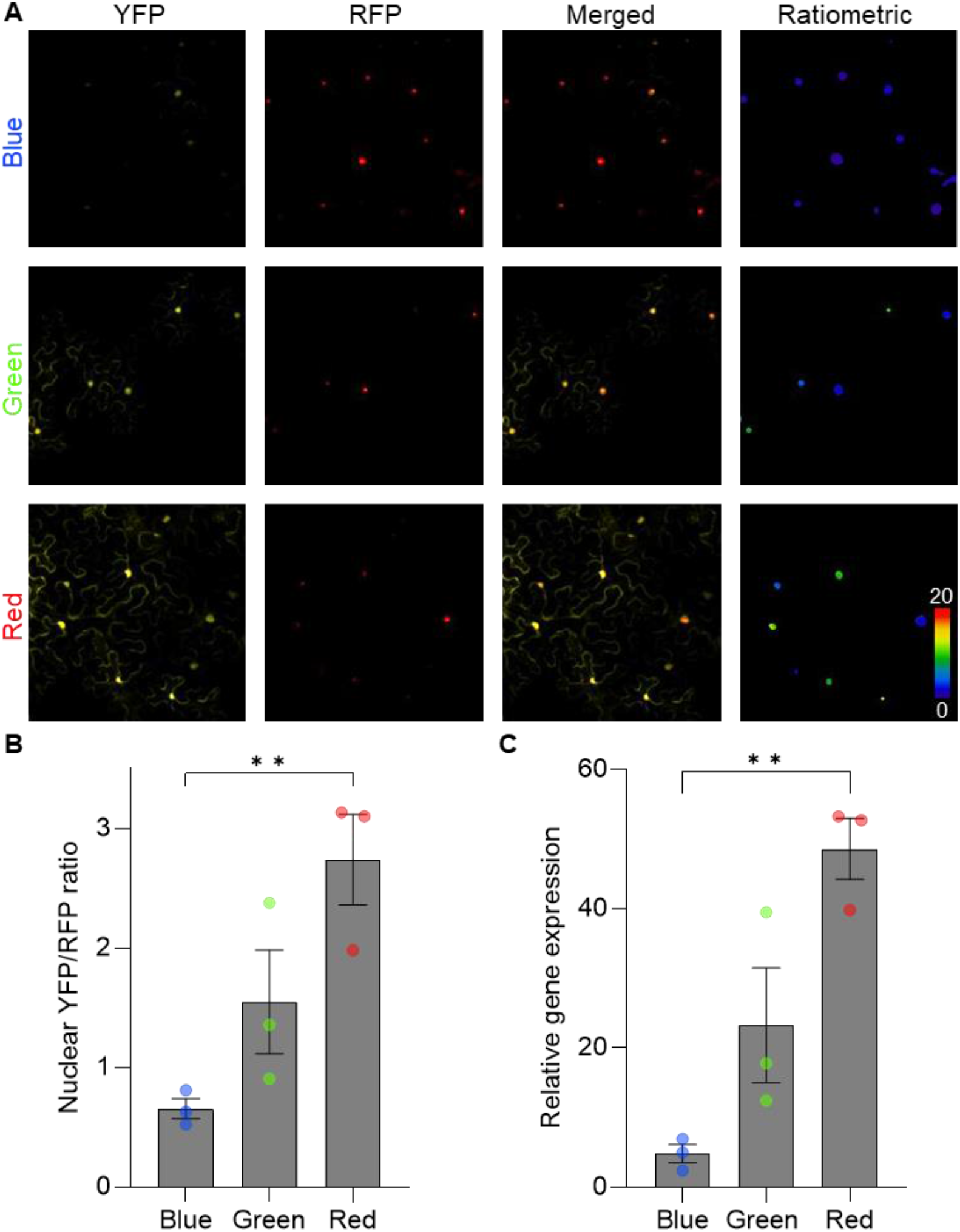
Deployment of Highlighter in *N. benthamiana* leaves for controlling fluorescent protein levels with monochromatic light. *N. benthamiana* leaves were infiltrated with Agrobacterium for delivery of Highlighter(YFP), and kept in darkness overnight before receiving continuous light treatments (100 μmol m^−2^ s^−1^) for 3 days with blue, green and red light from LEDs with peak wavelength emissions ∼ 455 nm, 525 nm and 660 nm, respectively. (A) Each column holds representative confocal images demonstrating nuclear YFP and RFP fluorescence in light treated samples, alongside merged images of the YFP and RFP fluorescence and finally the calculated YFP/RFP ratios (ratiometric). (B) Quantification of YFP/RFP ratios in light treated samples. (C) Relative gene expression levels in light treated samples. In (B) and (C), means and S.E.M. are presented for 3 biological independent experiments. Individual means are depicted with circles colored according to light treatment. **P<0.01; n per biological mean in (B) is 22 to 168 nuclei. Leaves were spot infiltrated with OD600 nm = 0.4 *A. tumefaciens* cultures.

### Optogenetic control of target gene expression with cellular resolution

Tunable system activity with cellular spatiotemporal resolution is a highly desirable property of optogenetic actuators and is required to answer a multitude of high-resolution biological hypotheses. To explore the spatiotemporal limits for gene induction with Highlighter, we locally irradiated neighboring regions in transiently transformed *N. benthamiana* leaf discs with 442 nm and 633 nm lasers using a Fluorescence recovery after photobleaching (FRAP) module on a confocal microscope. In leaf discs transformed with Highlighter(YFP), the YFP/RFP ratios remained low in the region treated with the 442 nm laser and steadily increased over time in the region treated with the 633 nm laser (Fig 4A and 4C). Transcript levels also correlated with this trend when exposed to similar red and blue stimuli generated with LED lights (Fig 4D). To investigate if the ratio-change observed was a consequence of photobleaching of the RFP control, we repeated the experiment using a response module variant where P_HL_ was exchanged with a 35S promoter variant for constitutive YFP expression. We did not observe substantial ratio changes in leaf discs when YFP was constitutively expressed (Fig 4B and 4C). To our knowledge, Highlighter therefore provides the first proof-of-concept for cellular expression control in planta using light stimuli in the visible spectrum.

**Fig 4.**
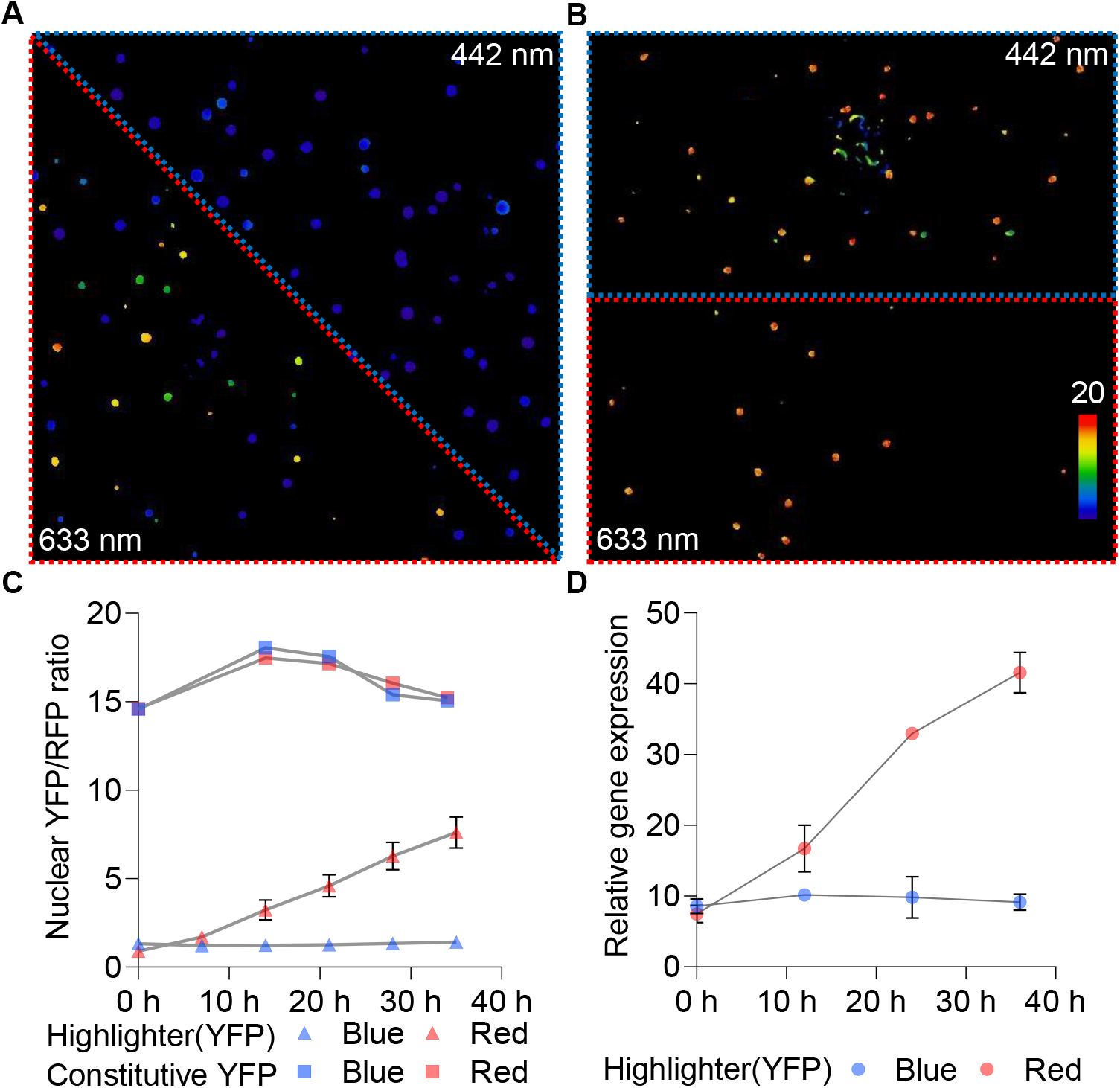
High-resolution control of target gene induction in *N. benthamiana* leaves using laser illumination. After infiltration with Agrobacterium for delivery of Highlighter(YFP) (A) or the constitutive YFP expression control (B), samples were kept in the dark overnight prior to continuously blue light treatment with LEDs (100 μmol m^−2^ s^−1^). Samples were blue light treated until 2.5 days post infiltration to minimize target gene expression and then subjected to blue and red light treatments with lasers (initiation of laser treatment are defined as 0 h in (C)). 442 nm blue and 633 nm red lasers were used to irradiate the area outlined in blue and red, respectively. Images in (A) and (B) are ratiometric representations of the YFP/RFP ratios observed after 38 h of light treatment (five 7 h light treatments interrupted by confocal imaging). Images are sum projections of z-stacks spanning multiple cell layers. Cellular resolution measurements of nuclear YFP/RFP ratios for Highlighter(YFP) are available in S4 Fig. (C) Temporal quantification of YFP/RFP ratios for laser-based target gene induction in (A) and (B). Highlighter(YFP) data is represented with triangles and data for constitutive YPF expression is represented with squares. Mean and S.E.Ms are presented for each time point. (D) Temporal quantification of YFP target gene transcript levels during blue or red light treatments. Again, samples were kept in the dark overnight and treated continuously with blue light until 2.5 days post infiltration (here defined as 0 h) to minimize target gene expression levels. Infiltrated leaves were then exposed to blue and red LED light treatments (λ ∼ 455 nm and 660 nm, respectively, 100 μmol m^−2^ s^−1^) for 36 h and leaf tissue was sampled every 12 h. Means and S.E.M. are presented for 3 biological independent experiments. Leaves were spot infiltrated with OD600 nm = 0.4 *A. tumefaciens* cultures.

### Optogenetic control of plant immunity

Highlighter was developed to assert control over biological processes *in planta*. Having demonstrated that Highlighter could be deployed to regulate target gene expression levels with cellular resolution, we aimed to provide proof-of-concept for application by modulating plant immune responses. Specifically, we tested whether Highlighter could be used for optogenetic control of the hypersensitive response (HR), a suite of responses activating effector-triggered immunity in plants [41]. High-resolution optogenetic control of HR in transiently transformed *N. benthamiana* would enable future experiments with sufficient spatiotemporal resolution for investigating the mechanisms underlying HR progression during infection.

For this demonstration, we chose to take optogenetic control over effector-triggered immunity with Highlighter by modulating expression levels of an auto-active immune regulatory protein. During effector-triggered immunity, plant cells recognize pathogen effector proteins with intracellular immune receptors called NLRs (nucleotide-binding and leucine-rich repeat). NLRs trigger innate immune responses, including rapid programmed cell death and accumulation of phenolic compounds in the local region surrounding an infection. Many sensor NLRs rely on helper NLRs, such as NLR-REQUIRED FOR CELL DEATH (NRC) proteins, to effectively translate effector recognition into HR [42]. A D478V mutation in NRC4 from *N. benthamiana* (NRC4^D478V^) creates and auto-active protein that can activate HR in the absence of infection when transiently overexpressed in *N. benthamiana* leaves [42]. NRC4^D478V^ thus presented an excellent target for Highlighter control of plant immunity.

We constructed Highlighter(NRC4^D478V^), as well as positive and negative controls, to regulate and evaluate effects of NRC4^D478V^ expression in *N. benthamiana* leaves (Fig 5A). To verify that constitutive expression of NRC4^D478V^ activates HR, we tracked the buildup of fluorescent compounds [43] during *Agrobacterium*-mediated transient expression. Strong, localized fluorescence was observed and readily imaged upon UV-A excitation (Fig 5B). The same was not observed for the negative control construct, which lacks the response regulator, CcaR_HL_ (Fig 5B). This suggested that the resulting fluorescence buildup could be used as proxy for NRC4^D478V^ expression and HR induction. For quantification of HR associated fluorescence in all experiments, negative control samples were used for background fluorescence subtraction while positive control samples were used for fluorescence normalization. During *Agrobacterium*-mediated transient expression of Highlighter(NRC4^D478V^), we observed markedly lower relative fluorescence in *N. benthamiana* leaves kept in continuous blue light (λ ∼ 455 nm), compared to green light (λ ∼ 525 nm), orange light (λ ∼ 630 nm) and red light (λ ∼ 660 nm) treatments (Fig 5B, and 6A). These results were in clear agreement with our previous results for optogenetic control of YFP/RFP ratios in transiently transformed *N. benthamiana* leaves (Fig 3).

**Fig 5.**
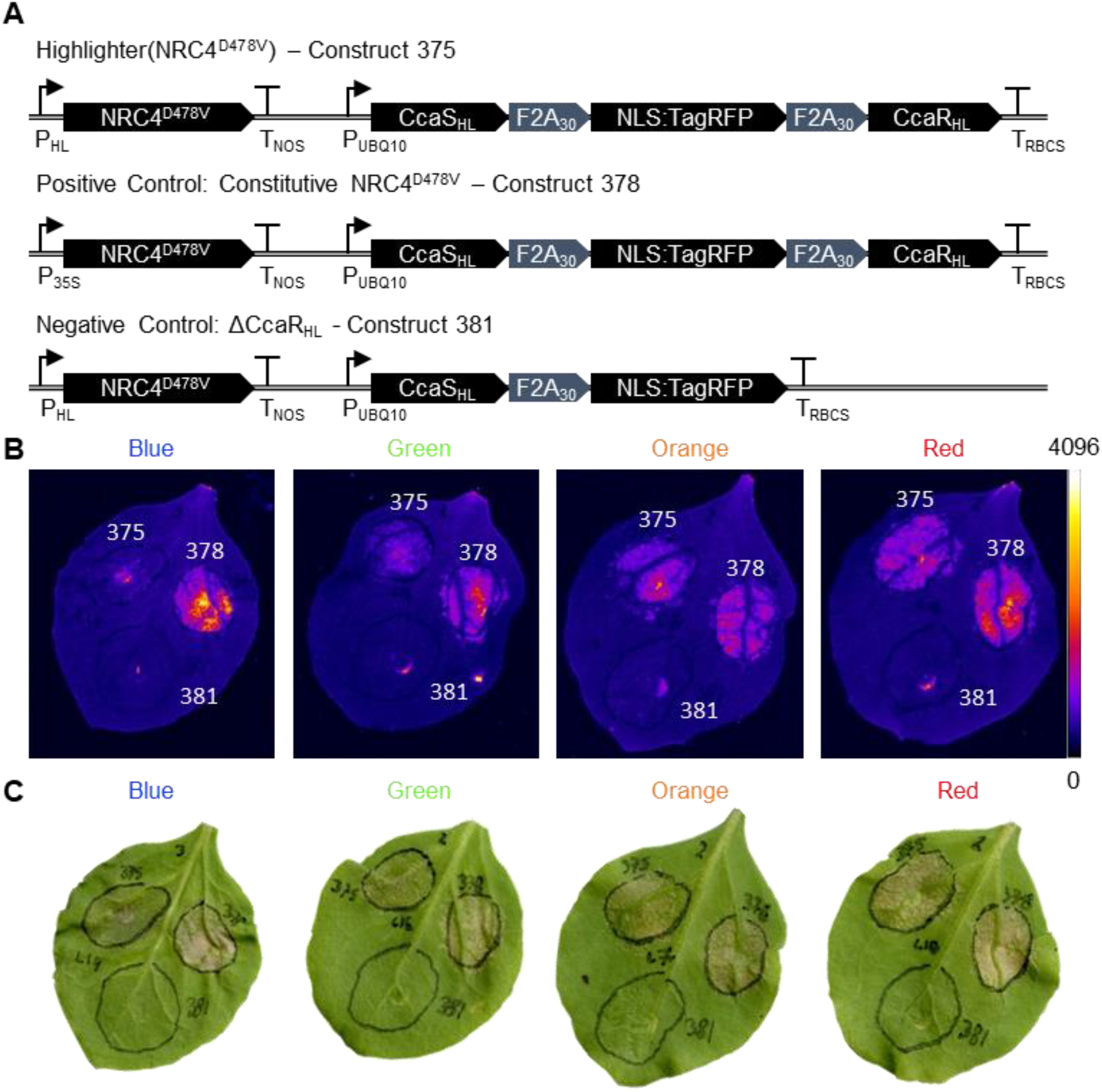
Highlighter controlled immune responses in transiently transformed *N. benthamiana* leaves. (A) Highlighter constructs used to assert control over immune responses in *N. benthamiana* leaves in response to light treatment. In Highlighter(NRC4^D478V^), construct 375, NRC4^D478V^ is under control of the Highlighter system, via P_HL_, whereas the Highlighter construct for constitutive NRC4^D478V^ expression, construct 378, is a positive control used for normalization. The Highlighter construct missing CcaR_HL_, construct 381, is a negative control construct used for background subtraction. (B) Representative images of HR-induced fluorescence in response to NRC4^D478V^ expression under blue, green, orange and red light; λ ∼ 455 nm, 525 nm, 630 nm and 660 nm, respectively. LUT is the Fire LUT, ImageJ. Strong UV-fluorescent signals are observed at the center of Agrobacterium-infiltrated spots due to tissue damage from the syringe infiltration and is also clearly observed in (C). (C) Images of leaves in (B) for demonstrating HR-associated cell death progression. Images in panels (B) and (C) were acquired 4 days post infiltration (DPI). Leaves were spot infiltrated with OD600 nm = 0.2 *A. tumefaciens* cultures.

**Fig 6.**
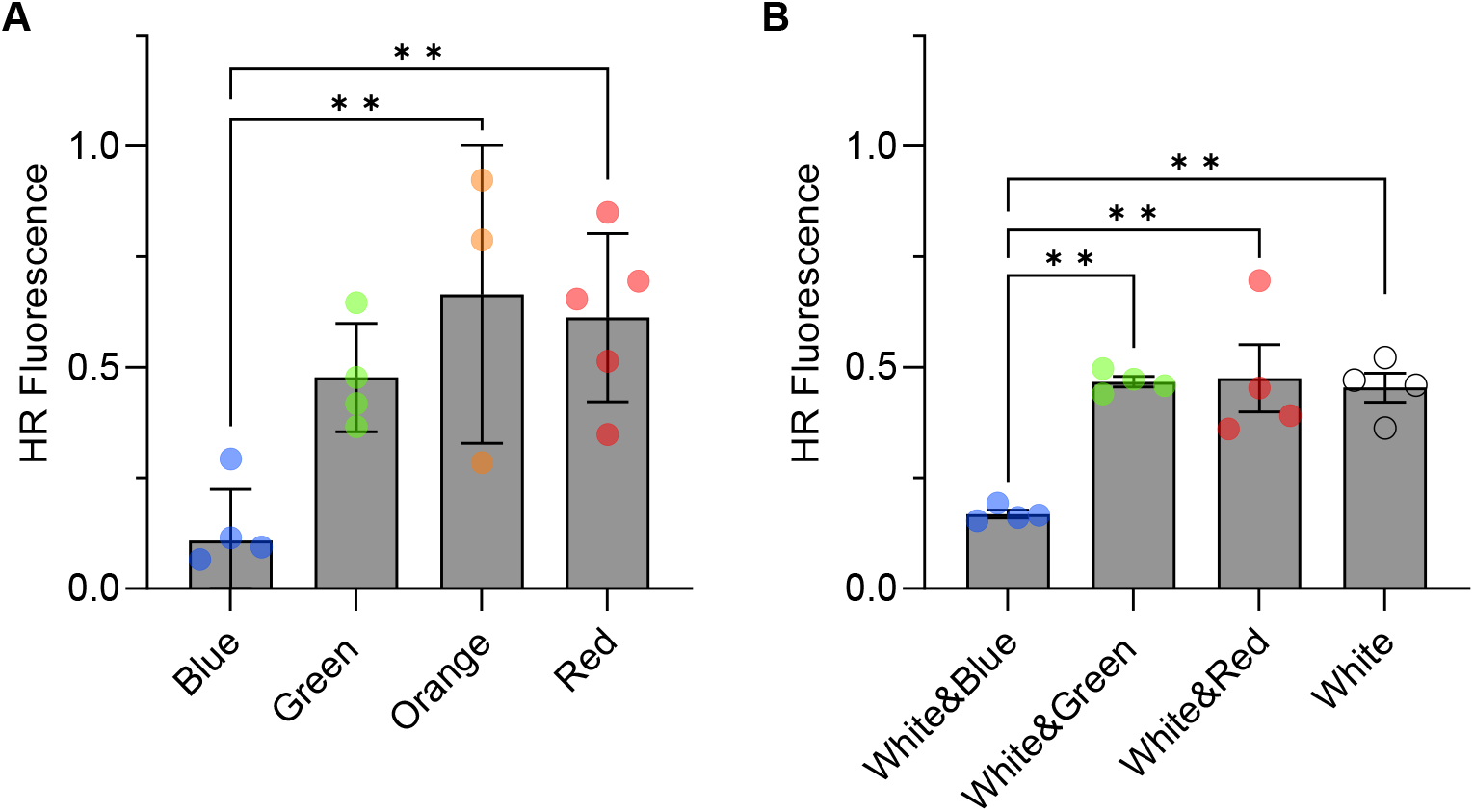
Highlighter controlled immune responses in *N. benthamiana* in response to monochromatic and modulated white light. (A) HR regulated by Highlighter-controlled NRC4 ^D478V^ expression under blue, green, orange and red LED light (λ ∼ 455 nm, 525 nm, 630 nm and 660 nm, 100 μmol m^−2^ s^−1^ light intensity). 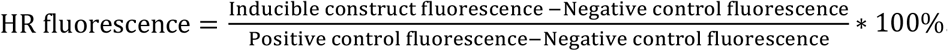. (B) Highlighter control of HR levels in white light regimes supplemented with blue, green or red light. Enriched white light regimes were defined as 50 μmol m^−2^ s^−1^ light from a 5700 K white light LED channel supplemented with 50 μmol m^−2^ s^−1^ light from blue, green, or red light channels with λ ∼ 455 nm, 525 nm and 660 nm, 100 μmol m^−2^ s^−1^ total light intensity. Mean and S.E.M are presented for each treatment and symbols/dots represent average HR responses from 3 to 5 biological repeats. n per biological average is 3 to 12 for monochromatic data in (A) and 5-14 for enriched white light data (B). * P<0.05 and ** P<0.01, statistics are One-way ANOVA. Leaves were spot infiltrated with OD600 nm = 0.2 *A. tumefaciens* cultures.

In standard horticultural environments, light typically originates from sunlight or complex broad-spectrum white light sources that support robust plant development. Consequently, to test if gene induction could be modulated under such conditions, we supplemented light from white light LEDs in a 1:1 ratio with blue, green and red light (total light intensity 100 μmol m^−2^ s^−1^). We observed low HR associated fluorescence in white light supplemented with blue light and high HR associated fluorescence in white light supplemented with green or red light (Fig 6B). This confirms that Highlighter is an optogenetic system that can control a biological process in plants grown in broad spectrum light.

## Discussion

The development of Highlighter, a cyanobacteriochrome-based light-inducible gene expression system for plants, represents an important step for high-resolution, minimally-invasive, and low-cost perturbation of plant biological processes. Advances in plant optogenetics have long been restricted by the limited availability of photoreceptor that are not native to plants and are independent of exogenously supplied chromophores. Ideal actuators in plants would also not photoswitch in horticultural light environments that include cycling between white light and darkness. Our conversion of the cyanobacterial CcaS-CcaR system for optogenetic control of target gene expression in plants is therefore an important innovation. The Highlighter technology exemplifies how spectrally diverse cyanobacteriochrome-based systems can be repurposed for optogenetic regulation of biological processes in plants, opening up a spectrum of new possibilities.

The complexity of light spectra and light-dark cycling in standard horticultural environments was until recently a fundamental challenge to contend with for optogenetic systems in plants. However, the PULSE system [16] elegantly demonstrated that this complication could be circumvented by combining two gene-expression switches with competing properties. PULSE combines an SRDX-EL222 “blue-off” module, to keep background gene expression low during the light cycle, and a PHYB-PIF “red-on” module, which is activated by monochromatic red light treatment. The Highlighter system, however, has an inherent “blue-off” response, without the need for an additional co-expressed module and thus makes it a simpler system for deployment in plants. The unexpected ‘blue-off’ response of the Highlighter system, however, warrants a follow-up investigation to determine its molecular basis in an inherently green/red sensor such as CcaS. Though unforeseen, inactivity in response to blue light is not unprecedented for a CcaS protein. The CcaS homolog from *Nostoc punctiforme* also demonstrated blue-off behavior when repurposed as an optogenetic actuator in *E. coli* [44]. For CcaS_HL_, our studies suggest that this response potentially arises from the blue-light mediated activity of a second CcaS-associated pigment, a flavin.

Unlike optogenetic systems based on plant photoreceptors, Highlighter is cyanobacterial in origin. This theoretically reduces the risk of Highlighter causing undesired off-target phenotypic effects and equally of endogenous light signaling pathways interfering with Highlighter activity. In the future, it will be interesting to test this in stable transgenic lines expressing Highlighter.

With advances in high-resolution quantitation, new hypotheses arise that can only be addressed by perturbing the measured biological process in precisely defined spatial regions and temporal windows. Such studies are often not feasibly conducted using chemically inducible systems because inducer molecules cannot be applied with sufficient resolution. Although future work will address these goals for Highlighter in stable transgenics, our transient expression studies, asserting optogenetic control over fluorescent reporter proteins and plant immunity, demonstrate that Highlighter is already a useful technology that allows precise optogenetic control of target gene expression down to the cellular level and can be deployed to modulate biological processes - even in complex light environments. From our experience using FRET biosensors to investigate how cellular hormone dynamics serve as signal integrators and major regulators of physiology and development [45,46], we also recognized a need to precisely perturb cellular hormone dynamics. We set out to develop Highlighter because we envisioned deploying the technology to evaluate hypotheses stemming from high-resolution measurements, for example distinguishing correlation from causation when investigating the connection between cellular gene expression or metabolite levels, and physiology and development. Beyond the scope of studying endogenous processes, the Highlighter technology holds great potential for plant biotechnology. Highlighter could address bottlenecks in transient *N. benthamiana*-based expression platforms for synthesis of high-value compounds and be used to optimally time developmental transitions or stress responses, such as immune activation to ward off pathogen outbreaks, in greenhouse or vertically farmed crops. We therefore expect Highlighter to become a resource in the optogenetic toolbox, changing how we approach hypothesis testing in plant biology and how we address production and yield bottlenecks in plant biotechnology.

## Materials and Methods

A detailed description of the plasmids used in this article, and their assembly, is found in S1 Table. PCR primers were synthesized by Sigma Aldrich (S2 Table), longer DNA fragments and genes were ordered from GeneScript (S3 Table). PCRs were performed using Q5 High-Fidelity DNA Polymerase (New England Biolabs (NEB), Cat#M0491S/L) and gel extractions were done with the Macherey-Nagel NucleoSpin Gel and PCR Clean-up Mini Kit (Macherey-Nagel, Cat#740609). DNA assemblies were carried out by In-Fusion Cloning (Takara Bio, In-Fusion HD Cloning Plus kit, Cat#638909) or NEBuilder assembly (NEB, NEBuilder High-Fidelity Master Mix, Cat#M5520) as per manufacturer’s instructions. Assembly reactions were transformed into chemically competent *E. coli* cells: Stellar competent cells (Takara Bio, Cat#636763), chemically competent DH5α cells or NEB 10-beta competent cells (NEB, Cat#C3019). Constructs were selected on LB plates (1% Tryptone, 0.5% Yeast Extract and 1% Sodium Chloride 1.5% Bacto agar) with appropriate selection. Plasmid purification was performed using the Qiagen QIAprep Spin Miniprep Kit (Qiagen, Cat#27106). Plasmids were verified by restriction enzyme digestion and sequencing (Sanger sequencing, Source BioScience). Site directed mutagenesis was performed using primers designed using the QuikChange Primer Design tool by Agilent Technologies with QuickChange® II Kit settings (https://www.agilent.com/store/primerDesignProgram.jsp).

*E. coli* strains were prepared for bacterial photoswitching experiments by co-transforming *E. coli* DH5α cells with vector sets for expressing the CcaS-CcaR system and system variants. One vector (based on pSR43.6r) expressed CcaS, or a CcaS variant, and genes for either PCB or PΦB biosynthesis, and a second vector (pBL413-003-020, derived from pSR58.6) expressed CcaR and further encoded an sfGFP reporter cassette where *sfgfp* is under the control of the engineered cognate promoter for CcaR, P_cpcG2-172_ [20]. Liquid *E. coli* cultures expressing CcaS-CcaR system variants were cultured in darkness for 12-14 h in LB (1% Tryptone, 0.5% Yeast Extract, 1% Sodium Chloride) with appropriate antibiotics in 96 well plates (VWR, Cat#732-3802), with one 3 mm glass bead and 750 μl media per well at 37°C, shaking at 220 rpm. Cultures were serial diluted in LB from 3-fold to 2187-fold in 96 well plates (Thermo Fisher Scientific, Greiner Bio-One™ Cat#655101) and incubated at 37°C, shaking (250 rpm) while receiving light treatments. Light treatments were ∼10 μmol m^−2^ s^−1^ light from LEDs with peak emissions around 400 nm, 455 nm, 525 nm, 590 nm, 605 nm, 630 nm, 660 nm and 695 nm. Complete spectra & LED models are found in S5 Fig. Light intensities were measured using a Licor LI-250A light meter with the LI-190R Quantum Sensor and spectra were recorded using an UPRtek MK350S LED meter. sfGFP fluorescence was quantified on a fluorimeter (Molecular devices, SpectraMax i3x - fluorescence read with 5 points, 6 flashes per well, bottom read, excitation 485 nm ± 4.5 nm and emission 516 nm ± 7.5 nm), along with the cell density (absorbance read at 600 nm, endpoint). For quantification of induction, fluorescence counts (LB fluorescence background subtracted) were plotted against cell densities (LB absorbance background subtracted). Fluorescence at OD600 nm = 0.2 was estimated from the plots with 3^rd^ order polynomial trendlines.

For heterologous expression, purification and spectroscopy of holo-CcaS_HL_, *E. coli* expressing hexahistidine tagged CcaS_HL_ alongside PΦB biosynthetic enzymes HO1 and mHY2 were cultured in 24 L of LB medium at 18°C. The purification protocol consisted of three steps performed on an ÄKTA Pure System (Cytiva): immobilized metal-affinity chromatography (Cytiva, 5 mL HisTrap HP, Cat#17524802) for hexahistidine tagged CcaS_HL_, ion-exchange chromatography with a linear gradient of salt (Cytiva, 5 mL HiTrap Q HP, Cat#17115401) and size-exclusion chromatography (SEC) (Cytiva, HiLoad 26/600 Superdex 200 pg, Cat#28989336). This expression and purification protocol yielded 3 mL of the target at ∼18 μM (determined from 280 nm absorbance of “post-purification” sample). Absorbance spectra (300-800 nm) for the resulting product was analyzed on a spectrometer (Agilent, Cary 60) following purification and then following illumination with light from the ColorDyne Benchtop Lightsource at the wavelengths and periods of time indicated in S2 Fig. In any given figure panel in S2 Fig, the same sample was illuminated for cumulative periods followed by data acquisition (e.g., illumination for 1 minute, followed by data acquisition (spectrum “1 min”); illumination for a further minute, followed by data acquisition (“2 min”); illumination for a further 3 minutes, followed by data acquisition (“5 min”); etc). Fluorescence emission spectra were acquired between 460-800 nm (S3 Fig) following photoexcitation at 445 nm of the red-absorbing state of CcaS_HL_ (to avoid further photoisomerisation of the PΦB chromophore during measurement).

*Agrobacterium*-mediated transient transformation and photoswitching assays in *N. benthamiana* were performed by transforming electrocompetent *A. tumefaciens* GV3101, carrying the pMP90 helper plasmid [47], with Highlighter plasmids in 1 mm electroporation cuvettes (Eurogentec, Cat#CE-0001-50) using an Eppendorf Multiporator (Cat#4308, 1500 V τ 5 ms). Cells recovered for 1-2 h in LB at 28°C and were selected on LB plates supplemented with appropriate antibiotics. *A. tumefaciens* strains carrying plasmids for testing Highlighter system variants in planta were cultured at 28°C in liquid LB media, shaking at 220 rpm, supplemented with appropriate antibiotics. Cultures were pelleted, washed and resuspended in infiltration media (10 mM MES, 10 mM MgCl_2_, 200 μM Acetosyringone (Sigma Aldrich, Cat#D134406), pH 5.6) to an OD600 nm of 0.2-4 and mixed equally with *A. tumefaciens* C58C1 cells carrying the p19 plasmid, encoding the p19 RNA-silencing suppressor from *Tomato bushy stunt virus* [48]. Four-week-old leaves were syringe infiltrated through the abaxial side and left in the dark for 8-16 h before undergoing light treatments. For light treatments, infiltrated leaves were cut from plants and placed on 1% water agarose plates, abaxial side up, and sealed with surgical tape. Light treatments of infiltrated leaves were performed using Heliospectra lamps (model RX30) with total light intensities of 100 μmol m^−2^ s^−1^. Monochromatic LED light regimes were generated using the blue light channel (450 nm), green light channel (530 nm), orange light channel (620 nm) and red light channel (660 nm). Enriched white light regimes were defined as 50 μmol m^−2^ s^−1^ light from 5700 K white light LEDs, enriched with 50 μmol m^−2^ s^−1^ light from one of the above mentioned blue, green and red LED channels. Light intensities were measured using a Licor LI-250A light meter with a LI-190R Quantum Sensor and spectra were recorded using an UPRtek MK350S LED meter (S7 Fig). HR responses were scored 4-5 days after infiltration via accumulation of HR associated fluorescent compounds in infiltrated spots using a Syngene G-BOX (Model F3-LFP, UV Transilluminator; manual capture mode, TLUM lighting, UV032 filter). UV-fluorescence response resulting from Highlighter induced NRC4^D478V^ expression was defined as follows: 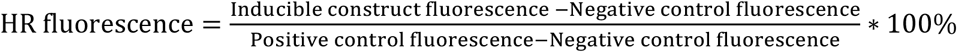. Fluorescent signals from nlsedAFPt9 and nlsTagRFP were collected by confocal imaging using a Leica TSC SP8 laser scanning confocal microscope. nlsedAFPt9 and nlsTagRFP were simultaneously excited with a 514 nm Argon laser; YFP emission was collected from 520-540 nm and RFP emission was collected from 595-625 nm on HyD detectors. Segmentation and quantification of fluorescence intensities were performed in ImageJ. 3D segmentation was performed using the 595-625 nm RFP channel and induction ratios were calculated as nuclear YFP signals divided by nuclear RFP signals. Overexposed voxels were excluded from the analysis when relevant. For high-resolution laser illumination to control target gene (nlsedAFPt9) expression levels, infiltrated plants were kept in darkness for 12-16 h post infiltration and continuously treated with blue light (100 μmol m^−2^ s^−1^ light from the 450 nm channel) until 2.5 days post infiltration. Infiltrated leaves were then cut off plants and transferred to 1% water agarose plates and placed under the objective on the confocal microscope. Cling film was used to seal the space between the plate and objective to maintain adequate humidity for sample health. A region with cells with early detectable nuclear localized RFP fluorescence was selected for time lapse imaging of nlsedAFPt9 expression. Light treatments were performed with a 442 nm laser (40 mW, 442 nm Diode laser at 0.45%) and a 633 nm laser (10 mW 633 nm HeNe laser at 0.15%) on a Leica TSC SP8 microscope using the FRAP module. Samples were light treated for 7h and imaged. Light treatment and imaging cycles were repeated up to five times.

For qRT-PCR quantification of gene expression levels, RNA was isolated from infiltrated, light-treated *N. benthamiana* leaf discs frozen in liquid nitrogen. Total RNA was extracted using the RNeasy Plant Mini Kit (Qiagen, Cat#74904) and DNase treated with the Invitrogen™ TURBO DNA-free™ Kit (Thermo Fisher Scientific, Cat#AM1907). cDNA was synthesized with the SuperScript VILO cDNA Synthesis Kit (Thermo Fisher Scientific, Cat#11754-250). Gene expression levels in samples were determined in quadruplicate by qPCR using gene-specific primers (5’GAAGAGAAAGGTTGGAGGGCT3’ and 5’TGACCGAAAACTTATGCCCGT3’ for nlsedAFPt9; 5’TGTGTCAGGGAAAGAATGGAG3’ and 5’TCAGAACCGAGCATATCGAG3’ for CcaR_HL_), a Lightcycler 480 (Roche Molecular Systems, Cat#05015243001) and qPCR LightCycler® 480 SYBR Green I Master (Roche Molecular Systems Cat# 04887352001) according to manufacturer’s instructions. Target gene (nlsedAFPt9) expression levels were quantified using the delta-delta Ct method [49], using CcaR_HL_ as the calibrator gene.

## Acknowledgements

The authors would like to thank Sebastian Schornack, James Locke, Tristan O. Kwan, Mike Shaw and Sophien Kamoun for providing feedback and Sophien Kamoun for sharing NRC4^D478V^ with us. We are grateful to Sophien Kamoun and Sebastian Schornack for inspiration and advice regarding asserting control over immune responses with Highlighter in *N. benthamiana* and to Temur Yunusov for providing us with technical advice and materials for transiently transforming N. benthamiana. James H. Rowe kindly provided technical assistance on image analysis. Also a special thanks to the late Winslow Russell Briggs for discussions and encouragement that helped us to initiate this project.

## Supporting information

**S1 Fig. Alignment of GAF domains from cyanobacteriochromes, plant phytochromes and bacterial phytochromes**.

**S2 Fig. Spectroscopic characterization of holo-CcaS**_**HL**_ **purified from PΦB-producing *E. coli*. S3 Fig. Fluorescence emission spectrum for holo-CcaS**_**HL**_ **purified from PΦB-producing *E. coli***.

**S4 Fig. Cellular resolution measurements of nuclear YFP/RFP ratios, from Fig 4C, generated by deploying Highlighter(YFP) in transiently transformed *N. benthamiana***.

**S5 Fig. Light spectra of LED arrays used for light treating *E. coli* cultures expressing the CcaS&CcaR system variants**.

**S6 Fig. Light spectra of LEDs used for the spectroscopic characterization of holo-CcaS**_**HL**_ **in S2 Fig. S7 Fig. Light spectra for Heliospectra RX30 lamps**.

**S1 Table. Vectors insert and construction description. S2 Table. Primers used to assemble vectors in S1 Table**.

**S3 Table. Synthesized genes used as PCR templates for vector assemblies**.

