## Supplementary material for "Highlighter: an optogenetic actuator for light-mediated, high resolution gene expression control in plants": S1 Table

| **S1 Table.** **Vectors insert and construction description.** | |
| --- | --- |
| Vector ID | Insert(s) |
| pSR43.6r | P_J23106 with sRBS (30000)_::*ccaS*:: T_rrnB T1_; P_J23108 with RBS pprotet_::*ho1::RBS pprotet::pcyA*:: T_rrnB T1_;  P_NEOKAN_::*specR*:T_Lambda_ _T0_ |
| pBL413-003-005 | P_J23106 with sRBS (30000)_::*ccaS*:: T_rrnB T1_; P_J23108 with RBS pprotet_::*ho1::RBS pprotet::mHY2*:: T_rrnB T1_;  P_NEOKAN_::*specR*:T_Lambda_ _T0_ |
| pBL413-006-058 | P_J23106 with sRBS (30000)_::*ccaS(A92V)*:: T_rrnB T1_; P_J23108 with RBS pprotet_::*ho1::RBS pprotet::mHY2*:: T_rrnB T1_; P_NEOKAN_:: *specR*:T_Lambda_ _T0_ |
| pBL413-020-201 | P_J23106 with sRBS (30000)_::*MM:nls:ccaS(del1_69 A92V):f2a30(del88_90)*:: T_rrnB T1_; P_J23108 with RBS pprotet_::*ho1::RBS pprotet::mHY2*:: T_rrnB T1_; P_NEOKAN_:: *specR*:T_Lambda_ |
| pBL413-003-020 | P_J23100 with sRBS (250)_::*ccaR*::T_rrnB T1_; P_cpcG2-172_::*sfgfp*:: T_rrnB T1_; P_Cat_:*kanR*::T_Lambda_ _T0_ |
| pBL413-024-257 | P_HL_::*nls:edAFPt9*::T_NOS_;  P_UBQ10_::*MM:nls:ccaS_CO4A_(del1_69 A92V):f2a_30_:nls:TagRFP:f2a_30_:ccaR_CO4A_:4xGSS:vp64_CD4A_:2xGGS:nls*::T_RBCS_;  P_NOS_::hygR::T_MAS_  Vector Backbone: LB T-DNA repeat, *oriV*, *trfA*, *ampR*, *ColEI* and RB T-DNA repeat |
| pBL413-024-259 | P_35S(-343 to -51)_::*nls:edAFPt9*::T_NOS_;  P_UBQ10_::*MM:nls:ccaS_CO4A_(del1_69 A92V):f2a_30_:nls:TagRFP:f2a_30_:ccaR_CO4A_:4xGSS:vp64_CD4A_:2xGGS:nls*::T_RBCS_;  P_NOS_::*hygR*::T_MAS_  Vector Backbone: LB T-DNA repeat, *oriV*, *trfA*, *ampR*, *ColEI* and RB T-DNA repeat |
| pBL413-024-260 | P_HL_::*nls:edAFPt9*::T_NOS_;  P_UBQ10_::*MM:nls:TagRFP:f2a_30_:ccaR_CO4A_:4xGSS:vp64_CD4A_:2xGGS:nls*::T_RBCS_;  P_NOS_::*hygR*::T_MAS_  Vector Backbone: LB T-DNA repeat, *oriV*, *trfA*, *ampR*, *ColEI* and RB T-DNA repeat |
| pBL413-024-261 | P_HL_::*nls:edAFPt9*::T_NOS_;  P_UBQ10_::MM:nls:ccaS_CO4A_(del1_69 A92V):f2a_30_:nls:TagRFP::T_RBCS_;  P_NOS_::*hygR*::T_MAS_  Vector Backbone: LB T-DNA repeat, *oriV*, *trfA*, *ampR*, *ColEI* and RB T-DNA repeat |
| pBL413-037-375 | P_HL_::*NRC4^D478V^*::T_NOS_;  P_UBQ10(-1094 to +370)_::*MM:nls:ccaS_CO4A_(del1_69 A92V):f2a_30_:nls:TagRFP:f2a_30_:ccaR_CO4A_:4xGSS:vp64_CD4A_:2xGGS:nls*::T_RBCS_;  P_NOS_::*hygR*::T_MAS_  Vector Backbone: LB T-DNA repeat, *specR*, *pBR322 Ori*, *bom*, *pVS1 oriV*, *pVS1 RepA*, *pVS1 StaA*, RB T-DNA repeat |
| pBL413-037-378 | P_35S(-343 to -51)_::*NRC4^D478V^*::T_NOS_;  P_UBQ10(-1094 to +370)_::*MM:nls:ccaS_CO4A_(del1_69 A92V):f2a_30_:nls:TagRFP:f2a_30_:ccaR_CO4A_:4xGSS:vp64_CD4A_:2xGGS:nls*::T_RBCS_; P_NOS_::*hygR*::T_MAS_  Vector Backbone: LB T-DNA repeat, *specR*, *pBR322 Ori*, *bom*, *pVS1 oriV*, *pVS1 RepA*, *pVS1 StaA*, RB T-DNA repeat |
| pBL413-037-381 | P_HL_::*NRC4^D478V^*::T_NOS_;  P_UBQ10(-1094 to +370)_::*MM:nls:ccaS_CO4A_(del1_69 A92V):f2a_30_:nls:TagRFP*::T_RBCS_;  P_NOS_::*hygR*::T_MAS_  Vector Backbone: LB T-DNA repeat, *specR*, *pBR322 Ori*, *bom*, *pVS1 oriV*, *pVS1 RepA*, *pVS1 StaA*, RB T-DNA repeat |
| pBL413-008-271 | P_T7_::*MM:6xH:nls:ccaS(del1_69 A92V):f2a30(del88_90)*::T_B0015_; P_J23108 with RBS pprotet_::*ho1::RBS pprotet::mHY2*:: T_rrnB T1_  P_NEOKAN_::*specR*:T_Lambda_ _T0_ |
| **Abbreviations**:  (BBa parts refer to parts from the Registry of Standard Biological Parts (<http://parts.igem.org/Main_Page)>)  2xGSS and 4xGSS: Two or four consecutive repeats of the Glycine-serine-serine linker.  sfgfp: Superfolder green fluorescent protein.  A92V: Mutation of amino acid alanine 92 in CcaS to valine.  *ampR*: CDS of the bla β-lactamase that confers resistance to ampicillin and carbenicillin.   - In the context of vector backbones (for pBL413-024-257, pBL413-024-259, pBL413-024-260 and pBL413-024-261), AmpR refers to the entire resistance gene and not simply the CDS.   bom: Basis of mobility region from pBR322.  CcaS: Chromatic acclimation sensor.  CcaR: Chromatic acclimation response regulator.  CD4A: This subscript annotation indicates that the nucleotide sequence has been codon-diversified to avoid identical repeat sequences and that codon usage is appropriate for expression in *A. thaliana.*  P_HL_: P_HL_ is a synthetic promoter comprised of 3 cis-regulatory elements, recognized by CcaR, fused to the 5’ end of the minimal plant promoter P_35Smin(-51)_.  CO4A: This subscript annotation indicates that the nucleotide sequence has been codon-optimized for expression in *A. thaliana.*  del: Deletion of bases from first numbered base to (_) last numbered base, both bases included.  edAFPt9: Enhanced dimerization variant of the yellow fluorescent protein Aphrodite, with a nine amino acid truncation. Aphrodite is a codon-modified version of Venus.  ColEI: High-copy-number ColE1 origin of replication.  F2A30: Long F2A ribosomal skipping sequence variant, comprising 30 codons, adding 29 amino acids to the C-terminus of the upstream protein and a proline residue on the N-terminus of the downstream protein during translation.  *ho1*: Heme oxygenase 1 gene.  HygR: CDS of the *hph* aminoglycoside phosphotransferases gene from *E. coli* that confers resistance to hygromycin.  KanR: CDS for an aminoglycoside phosphotransferase that confers resistance to Kanamycin in bacteria.  LB T-DNA repeat: T-DNA left border repeat from the nopaline Ti plasmid C58.  M: ATG codon for methionine.  *mHY2*: gene encoding the mHY2 PΦB synthase from *Arabidopsis* without the transit peptide.  *nls*: Nuclear localization sequence.  NRC4^D478V^: Autoactive helper NLR (nucleotide-binding and leucine-rich repeat domains) protein from *N. benthamiana* with the D478V mutation.  oriV: incP origin of replication.  P: Promoter (Promoter name/description in subscript).   - Constitutive bacterial promoter P_J23100_, P_J23106_ and P_J23108_ are parts BBa_J23100, BBa_J23106, and BBa_J23108, respectively. - Constitutive bacterial promoter P_NEOKAN_ is from the neomycin-kanamycin resistance gene. - P_cpcG2-172_ is a truncated cognate cyanobacterial promoter for CcaR. - P_Cat_ is a bacterial promoter from the chloramphenicol acetyltransferase gene. - P_NOS_ is a constitutive plant promoter for the nopaline synthase gene. - P_35S(-343 to -51)_ is a truncated variant of the constitutive Cauliflower Mosaic Virus-35S promoter - P_UBQ10_ is the constitutive polyubiquitin 10 promoter from *Arabidopsis*.   pBR322 Ori: High-copy-number pBR322 origin of replication.  *pcyA*: Ferredoxin-dependent bilin reductase gene.  pVS1 oriV: Origin of replication for the Pseudomonas plasmid pVS1 (Heeb et al., 2000).  pVS1 StaA: Stability protein from the Pseudomonas plasmid pVS1 (Heeb et al., 2000).  pVS1 RepA: Replication protein from the Pseudomonas plasmid pVS1 (Heeb et al., 2000).  RB T-DNA repeat: T-DNA right border repeat from the nopaline Ti plasmid C58.  RBS: Ribosomal binding site.   - sRBS indicates it is a strong RBS. - sRBS (250), sRBS(30000) and RBS pprotet are parts BBa_K3165004, BBa_K3165005 and BBa_B0034, respectively.   SpecR: CDS for SmR, an aminoglycoside adenylyltransferase, aadA , that confers resistance to spectinomycin and streptomycin.   - In the context of vector backbones (for pBL413-037-375, pBL413-037-378 and pBL413-037-381), SpecR refers to the entire resistance gene and not simply the CDS.   T: Terminator (Terminator name/description in subscript).   - Bacterial terminator T_rrnB T1_ is part BBa_B0015, a double terminator made from transcription terminator T1 from the *E. coli* rrnB gene and early transcription terminator from phage T7. - Bacterial terminator T_Lambda_ _T0_ is part BBa_K1897030, the T0 transcription terminator from phage lambda. - T_NOS_ is the plant terminator and poly(A) signal for the nopaline synthase gene. - T_RBCS_ is the plant terminator from the Rubisco small subunit 1A gene from *Arabidopsis*. - T_MAS_ is the plant terminator from the mannopine synthase gene.   TagRFP: Monomeric red/orange fluorescent protein made from red fluorescent protein from *Entacmaea quadricolor* (Merzlyak et al., 2007).  VP64: Tetrameric VP16 transcription activator domain from human cytomegalovirus (Seipel K, 1992).  trfA: Trans-acting replication protein that binds to and activates oriV. | |
| **Description of cloning procedure**:  **pSR43.6r** was ordered from AddGene  **pBL413-003-005** was constructed by replacing the *pcyA* CDS with the *mHY2* CDS. *mHY*2 was amplified from *HY2*, a PΦB synthase from *Arabidopsis* (ordered as a cDNA clone from ABRC - clone ID U25336, AT3G09150 cDNA). The *mHY2* gene fragment was amplified with primers BL413-003-P001 and BL413-003-002. This fragment was used to replace *pcyA* in pSR43.6r in a three fragment In-Fusion HD cloning assembly. The pSR43.6r vector, except for the *pcyA* gene, was amplified as two PCR fragments with primers BL413-003-P003 and BL413-003-P015 and with BL413-003-P004 and BL413-003-P005.  **pBL413-006-058** was made from pBL413-003-005 by site directed mutagenesis with primers BL413-006-P099 and BL413-006-P100.  **pBL413-020-201** was made sequentially. First, the A92V mutation was introduced into CcaS in pSR43.6r by site directed mutagenesis with primers BL413-006-P099 and BL413-006-P100 to generate pBL413-006-050. Second, the N-terminal transmembrane domain of CcaS(A92V) was replaced with a nuclear localization sequence and *pcyA* was replaced with *mHY2* for phytochromobilin synthesis. This was done by In-Fusion HD cloning where the vector backbone, including *mHY2*, of pBL413-003-005 was amplified with BL413-003-P051 and BL413-004-P128 and the transmembrane domain in CcaS(A92V) was replaced with an NLS, using primers BL413-004-P127 and BL413-003-P050 with pBL413-006-050 as template. The resulting vector pBL413-004-135 was finally used in a 3 fragment In-Fusion HD cloning assembly to generate pBL413-020-201 by introducing an additional start codon 5’ of the nls:CcaS(A92V) CDS and adding the 5’ 87 nucleotides of the F2A_30_ sequence to the 3’ end of the nls:CcaS(A92V) CDS. Primer pairs used for these modifications were BL413-020-P321 and BL413-020-P328, BL413-020-P327 and BL413-003-P023, and BL413-003-P022 and BL413-020-P322.  **pBL413-003-020** was constructed from pSR58.6 (ordered from AddGene) by replacing the vector’s chloramphenicol resistance gene with the kanamycin resistance gene from pDONR221. The Kanamycin resistance gene was amplified with primers BL413-003-P091 and BL413-003-P092. All of pSR58.6, except for the chloramphenicol resistance gene was amplified using primers BL413-003-P093 and BL413-003-P094.  **pBL413-024-257** was made sequentially, starting with pUBQ-USER-2, a plant expression vector based on pEAQ, which was obtained from Meike Burow (Department of Plant and Environmental Sciences, Faculty of Science, University of Copenhagen). First, the KanR CDS was replaced in pUBQ-USER-2 with the AmpR CDS from pOO2 (obtained from Wolf Frommer, Carnegie institution for Science). The AmpR CDS was amplified with primers BL413-012-P138 and BL413-012-P139 from pOO2 and the whole pUBQ-USER-2 vector, except for the KanR CDS, was amplified with primers BL413-012-P140 and BL413-012-P141 and assembled by In-Fusion HD cloning to create pBL413-012-148. Second a cassette for target gene expression was entered into pBL413-012-148 by In-Fusion HD cloning to create pBL413-012-151. The cassette was composed of the bacterial cognate promoter for CcaR, P_cpcG2-172_, a minimal promoter (P_35Smin(-51)_), a gateway cassette with attR1 and attR2 (but no cmR and ccdB genes ) and a terminator (T_NOS_). P_cpcG2-172_ was amplified from pSR58.6 with primers BL413-012-P142 and BL413-012-P143; P_35Smin(-51)_ and attR1 was amplified from pMDC7 with primers BL413-012-P174 and BL413-012-P181; attR2 was amplified from pMDC7 with primers BL413-012-P180 and BL413-012-P182; T_NOS_ was amplified from pPTHyg with primers BL413-012-P147 and BL413-012-P148. To reduce the complexity of the assembly, the “P_cpcG2-172_“ fragment and the “P_35Smin(-51)_ and attR1” fragment was combined by overlap PCR (using BL413-012-P142 and BL413-012-P181) and the “attR2” fragment was combined with the “T_NOS_“ fragment (using BL413-012-P180 and BL413-012-P148). These two resulting fragments were introduced into the pBL413-012-148 vector upstream of the UBQ10 promoter. The pBL413-012-148 backbone was amplified as two fragments with primers BL413-012-P144 and BL413-012-P170 and with BL413-012-P149 and BL413-012-P171. Third, CcaS_HL_ and CcaR were codon-optimized for *A. thaliana* (CO4A) and entered into pUBQ-USER-2 to be expressed from the UBQ10 promoter-RBCS terminator cassette. To accomplish this, codon-optimized CDS templates for CcaS and CcaR (MM:NLS:CcaS(A92V del1_87) and CcaR MM:NLS:CcaR:VP16) were synthesized for the assembly by Genscript, and a P2A ribosomal skipping sequence (Khosla, 2020) would be introduced between CcaS_HL_ and CcaR in primer overhangs for transcriptional and translational linkage. Codon-optimized CcaSHL and NLS:CcaR(del1_3)VP16 was amplified with primers BL413-012-P187 and BL413-012-P188, and with BL413-012-P189 and BL413-012-P190, respectively, and the pUBQ-USER-2 backbone was amplified as two fragments using primers BL413-012-P183 and BL413-012-P184, and primers BL413-012-P185 and BL413-012-P186. The four fragments were assembled by In-Fusion HD cloning to create pBL413-012-152. (BL413-012-P186 and BL413-012-P187 introduced a double start codon and NLS to the 5’ end of the codon-optimized CcaS CDS amplified from MM:NLS:CcaS(A92V del1_87) and reintroduced bases 70 to 87 in CcaS). Fourth, the target gene expression cassette (P_cpcG2-172_:P_35Smin(-51)_::attR1 attR2:: T_NOS_) from pBL413-012-151 and the system expression cassette (P_UBQ10_::MM:nls:ccaS_CO4A_(del1_69 A92V):P2A_CO4A_:ccaR_CO4A_::T_RBCS_) from pBL413-012-152 were combined into a single vector in a three fragment In-Fusion HD cloning assembly creating pBL413-012-156. The target gene expression cassette from pBL413-012-151 was amplified with primers BL413-012-P142 and BL413-012-P177, while the system expression cassette and accompanying vector backbone was amplified from pBL413-012-152 with primers BL413-012-P149 and BL413-012-P171 and with BL413-012-P144 and BL413-012-P170. Fifth, the KanR CDS was replaced in pBL413-012-156 with the AmpR CDS from pBL413-012-156, to create pBL413-012-160, as described above for pBL413-012-148, but using primer BL413-012-P186 with BL413-012-140 and BL413-012-P187 with BL413-012-141 to amplify the vector backbone of pBL413-012-156 in two fragments. Sixth, the CmR and ccdB genes were then introduced between attR1 and attR2 to complete the gateway cassette, creating pBL413-012-150. This was achieved by restriction ligation cloning using the restriction enzyme SalI and pMDC7 as the donor vector for the CmR and ccdB genes. Seventh, nls:edAFPt9 was entered into pBL413-012-150 as the target gene. attB1 nls:edAFPt9 attB2 was amplified from pFLIPnls43 with primers BL413-012-P235 and BL413-012-P236 and nls:edAFPt9 was entered into pDON221, to create pBL413-012-162 in a Gateway BP reaction. nls:edAFPt9 was subsequently transferred to pBL413-012-150 in a gateway LR reaction, creating pBL413-012-164. Eight, F2A_30_:TagRFP was added N-terminally to MM:nls:ccaS_CO4A_(del1_69 A92V), creating pBL413-018-181 from pBL413-012-164 in a four fragment In-Fusion HD cloning assembly. TagRFP was amplified from the 2R3e_4xgly_TagRFP vector (provided by the Helariutta Group, SLCU, Cambridge University, UK) with primers BL413-020-P256 and BL413-020-P267. pBL413-012-164 was amplified in three fragments using primers BL413-020-P266 and BL413-020-P252 and with BL413-020-P251 and BL413-020-P253 and with BL413-020-P247 and BL413-020-P255. The F2A30 ribosomal skipping sequence was added via the overhangs of BL413-020-P255 and BL413-020-P256. Ninth, ccaR, codon-optimized for *Arabidopsis*, with a 5’ F2A30 sequence and 3’ nuclear localization signal and VP40 transcription activation domain was fused to the c-terminal end of TagRFP in pBL413-018-181 in a 4 fragment In-Fusion HD cloning assembly. The nuclear localization signal and VP40 transcription activation domain was ordered synthetically from Genewiz and was amplified using primers BL413-021-P363 and BL413-021-P364. Codon-optimized ccaR was amplified from pBL413-012-164 with primers BL413-021-P361 and BL413-021-P365. pBL413-018-181 was amplified, without the stop codon on TagRFP, in two fragments using primers BL413-021-P366 and BL413-019-P246 and with BL413-019-P245 and BL413-019-P362. In the assembly, the overhangs of BL413-019-P245 and BL413-019-P246 additionally replaced P_cpcG2-172_ with three cis-regulatory elements for CcaR, that together with the 3’ P_35Smin(-51_ sequence constitutes P_HL_. Tenth, to prepare Highlighter for stable transformation of plants, at a later stage in the project, a hygromycin selection cassette was assembled in the pUC19 vector before moving the selection cassette into pBL413-021-213. The NOS promoter and Hygromycin CDS from pPTHyg was amplified with primers BL413-024-P382 and BL413-024-P383, the MAS terminator was amplified from p16TKan with primers BL413-024-P380 and BL413-024-P381 and the pUC vector backbone was amplified with primers BL413-024-P378 and BL413-024-P379. The three fragments were assembled by In-Fusion HD cloning, creating pBL413-024-232. The HygR resistance cassette from pBL413-024-232 was transferred to pBL413-021-213 in a 4 fragment NEuilder assembly. The HygR resistance cassette was amplified with primers BL413-024-P388 and BL413-024-P389. pBL413-021-213 was amplified in three fragments with primers BL413-024-P390 and BL413-024-P391, with BL413-024-P392 and BL413-024-P387 and with BL413-024-P386 and BL413-024-P393. BL413-024-P386 and BL413-024-P387 additionally introduced a NLS to TagRFP via the primers overhangs. The resulting vector from this assembly is pBL413-024-230. Finally to introduce a stop codon to the Hygromycin CDS cassette in pBL413-024-230, the vector was amplified with primers BL413-024-P392 and BL413-024-P429 and with BL413-024-P430 and BL413-024-P391, to create pBL413-024-257 in a 2 fragment NEBuilder assembly.  **pBL413-024-259** was assembled from pBL413-024-257 and pBL413-026-242. pBL413-026-242 holds a 35S(-343 to -51) promoter which was originally amplified from pPTHyg with primers BL413-026-P409 and BL413-026-P410. The 35S(-343 to -51) promoter was re-amplified with primers BL413-024-P459 and BL413-024-P460 to replace P_HL_ in pBL413-024-257 and the rest of pBL413-024-257 was amplified as two fragments with primers BL413-024-P461 and BL413-024-P429 and with BL413-024-P430 and BL413-024-P462. The three fragments were assembled into pBL413-024-259 in a three fragment NEBuilder assembly.  **pBL413-024-260** was assembled in a two fragment NEBuilder assembly by removing nls:ccaS_CO4A_(del1_69 A92V):f2a_30_ from pBL413-024-257. For this, pBL413-024-257 was amplified with primers BL413-024-P481 and BL413-024-P482 and with BL413-024-P483 and BL413-024-P484.  **pBL413-024-261** was assembled in a two fragment NEBuilder assembly by removing :f2a_30_:ccaR_CO4A_:4xGSS:vp64_CD4A_:2xGGS:nls from pBL413-024-257. For this, pBL413-024-257 was amplified with primers BL413-024-P485 and BL413-024-P391 and with BL413-024-P392 and BL413-024-P486.  **pBL413-037-375**: To create pBL413-037-375, the short UBQ 10 promoter (-264 to + 370) in pBL413-024-257 was first replaced with the longer UBQ10 promoter (-1094 to +370) from the pUB1500 destination GW vector (provided by Sebastian Schornack, SLCU, Cambridge University, UK). The promoter was amplified with primers BL413-027-P603 and BL413-027-P604. pBL413-024-257 was amplified in two fragments using primers BL413-027-P605 and BL413-027-P606 and using BL413-027-P607 and BL413-027-P608. The three fragments were assembled in a NEBuilder assembly to create pBL413-027-315. Second, the vector backbone, including left border and right border sequences, was replaced in pBL413-027-315 with that of p16Kan in a three fragment NEBuilder assembly, creating pBL413-037-366. The p16Kan vector backbone was amplified with primers BL413-037-P705 and BL413-037-P706 and everything inside the left and right borders in pBL413-027-315 was amplified with BL413-037-P707 and BL413-037-P681 and with BL413-008-P200 and BL413-037-P708. Third, the nls:edAFPt9 CDS in pBL413-037-366 was replace with the CmR and ccdB genes from pDONR221 in a BP gateway reaction. Fourth, to complete pBL413-037-375 for NRC4^D478V^ expression from p_HL_, NRC4^D478V^ was amplified from an *Agrobacterium tumefaciens* GV3101 strain previously transformed with a vector for NRC4^D478V^ expression. NRC4^D478V^ was amplified with primers BL413-037-P701 and BL413-037-P702. The PCR product was entered into pDONR221 in a two fragment NEBuilder assembly where the pDONR221 backbone was amplified with primers BL413-037-P697 and BL413-037-P698, creating pBL413-037-363. Fifth, NRC4^D478V^ was entered into pBL413-037-366 in a LR gateway reaction with pBL413-037-363 to create pBL413-037-375.  **pBL413-037-378** was assembled following the identical procedure as that used for pBL413-037-375, but pBL413-037-378 is based on pBL413-024-259 instead of pBL413-024-257.  **pBL413-037-381** was assembled following the identical procedure as that used for pBL413-037-375, but pBL413-037-381 is based on pBL413-024-261 instead of pBL413-024-257. | |
