## Supplementary material for "Highlighter: an optogenetic actuator for light-mediated, high resolution gene expression control in plants": S2 Table

| **S2 Table.** **Primers used to assemble vectors in S1 Table.** Sequences annealing to templates are underscored. | |
| --- | --- |
| Primer ID | Primer sequence |
| BL413-003-P001 | AAGGATCCATGAGAGTCTCTGCTGTGTCGTATAAG |
| BL413-003-P002 | GGTCTAGATTAGCCGATAAATTGTCCTGTTAAATCC |
| BL413-003-P003 | TCGGCTAATCTAGACCAGGCATCAAATAAAACGA |
| BL413-003-P004 | ACTCTCATGGATCCTTTCTCCTCTTTAACTAGCC |
| BL413-003-P005 | GGCCTCAAATACAAAGTCTGCA |
| BL413-003-P015 | TTTGTATTTGAGGCCAGACTCC |
| BL413-003-P022 | GTAAGGCTTGATGAAACAACG |
| BL413-003-P023 | TTCATCAAGCCTTACGGTC |
| BL413-003-P050 | GAACACCATAATGCCGATC |
| BL413-003-P051 | GGCATTATGGTGTTCAACAG |
| BL413-003-P091 | AAGCTAAAATGAGCCATATTCAACGG |
| BL413-003-P092 | ATATCAAATTAGAAAAACTCATCGAGCAT |
| BL413-003-P093 | TGGCTCATTTTAGCTTCCTTAGCTCCTG |
| BL413-003-P094 | TTTTCTAATTTGATATCGAGCTCGCT |
| BL413-004-P127 | AAGAAGAAAAGGAAGGTGGGTGGAAGACAAAACCAAGAACGCC |
| BL413-004-P128 | CCTTCCTTTTCTTCTTTGGTTGTAACATTGCGCCTTCCTCCTA |
| BL413-006-P099 | CGATTCCGTAATGACGCTGCCCGTGCC |
| BL413-006-P100 | GGCACGGGCAGCGTCATTACGGAATCG |
| BL413-008-P200 | ACGTTCGTCAAGTTCAATGCA |
| BL413-012-P138 | AATAATAAATGAGTATTCAACATTTCCGTG |
| BL413-012-P139 | TCTAGGTATTACCAATGCTTAATCAGTGAGG |
| BL413-012-P140 | ATACTCATTTATTATTTCCTTCCTCTTTTCTACAG |
| BL413-012-P141 | ATTGGTAATACCTAGATGTGGCGCAAC |
| BL413-012-P142 | CGCAACTGAGCCCATTGTGCTTTTCTCT |
| BL413-012-P143 | GAGTCGAGTTAAAAATGCGATCCTAACAAAG |
| BL413-012-P144 | AATGGGCTCAGTTGCGCAGCCTGAATGG |
| BL413-012-P147 | AAAGTGGTTAGAGTAGATGCCGACCGAACAAG |
| BL413-012-P148 | CTTCCCAAATCAGCTTGCATGCCGGT |
| BL413-012-P149 | AAGCTGATTTGGGAAGGGACCCGACGAG |
| BL413-012-P170 | GCAAACAGCACGACGATTTCCTC |
| BL413-012-P171 | TCGTCGTGCTGTTTGCTGGC |
| BL413-012-P174 | ATCGCATTTTTAACTCGACTCTAGGATCTTCGC |
| BL413-012-P177 | CTTCCCAAATCAGCTTGCAT |
| BL413-012-P180 | AAGCTAAAATGTCAGGCTCCCTTATACACAGC |
| BL413-012-P181 | CCTGACATTTTAGCTTCCTTAGCTCCTGA |
| BL413-012-P182 | GGTCGGCATCTACTCTAACCACTTTGTACAAGAAAGCTGAACG |
| BL413-012-P183 | GAAAGAATTGATAAATTAAACCTCAGCAGCTTTCG |
| BL413-012-P184 | GAGAAGTACCGCAAGCTGTC |
| BL413-012-P185 | GCTTGCGGTACTTCTCCC |
| BL413-012-P186 | CCACCCACCTTCCTTTTCTTCTTTGGTTGTAACATCATATTAAGCCTCAGCGATCCC |
| BL413-012-P187 | AAAGGAAGGTGGGTGGAAGACAGAACCAAGAACGAAGAAGAATAGAAATAAGTATCAAGCAGCAG |
| BL413-012-P188 | CCAGCTTGCTTGAGCAATGAGAAGTTAGTAGCTCCGCTTCCAGCCCTTGGGAGATGGTT |
| BL413-012-P189 | TGCTCAAGCAAGCTGGAGATGTGGAAGAAAATCCTGGACCTAGAATACTCCTCGTGGAAGATGA |
| BL413-012-P190 | ATTTATCAATTCTTTCCCTGACACAAAGACT |
| BL413-012-P235 | GGGGACAAGTTTGTACAAAAAAGCAGGCTATGCTGCAGCCTAAGAAGAAGAGAAAGGTTGGAGG |
| BL413-012-P236 | GGGGACCACTTTGTACAAGAAAGCTGGGTTCATATGCCCGCCGCCGTGAG |
| BL413-019-P245 | TCCGATTTCTTTACGATTTGGCTTTCCGATTTCTTTACGATTTATCCTTCGCAAGACCCTTCCT |
| BL413-019-P246 | TCGTAAAGAAATCGGAAAGCGGAAATCGTAAAGAAATCGGAAAGCCAGTTGCGCAGCCTGAATG |
| BL413-020-P247 | GAACTGAAGAGGTTAGACTTGCTTT |
| BL413-020-P251 | GGCGCTTTCTCATAGCTCA |
| BL413-020-P252 | GCTATGAGAAAGCGCCAC |
| BL413-020-P253 | CTAACCTCTTCAGTTCTTCTTCTCTCT |
| BL413-020-P255 | TTAGCAAGTCAAAGTTGAGAGTCTGCTTCACCGGTGCCACAATTTTCTGTTTGTGAGCCCTTGGGAGATGGTT |
| BL413-020-P256 | AACTTTGACTTGCTAAAGTTAGCTGGTGATGTTGAATCTAATCCTGGACCAAGCGAGCTGATTAAGGAGAACA |
| BL413-020-P266 | GGCACAAGTGATAAATTAAACCTCAGCAGCTT |
| BL413-020-P267 | ATTTATCACTTGTGCCCCAGTTTGCT |
| BL413-020-P321 | GCGCAATGATGTTACAACCAAAGAAGAAAAGG |
| BL413-020-P322 | TGTAACATCATTGCGCCTTCCTCCTATTA |
| BL413-020-P327 | AACTTTGACTTGCTAAAGTTAGCTGGTGATGTTGAATCTAATCCTGGATAATAATCTAGACCAGGCATCAAAT |
| BL413-020-P328 | TAGCAAGTCAAAGTTGAGAGTCTGCTTCACCGGTGCCACAATTTTCTGTTTGTGAGCTCGAGGCAAATGGTTA |
| BL413-021-P361 | AACTTTGACTTGCTAAAGTTAGCTGGTGATGTTGAATCTAATCCTGGACCAATGAGAATACTCCTCGTGGAAGATGA |
| BL413-021-P362 | TTAGCAAGTCAAAGTTGAGAGTCTGCTTCACCGGTGCCACAATTTTCTGTTTGTGCTTGTGCCCCAGTTTGCT |
| BL413-021-P363 | GAAAGAATGGAGGCTCCGGTGGGTCA |
| BL413-021-P364 | TTTATCAACCTCCAACCTTTCTCTTCTTCTTAGGCT |
| BL413-021-P365 | AGCCTCCATTCTTTCCCTGACACAAAGACTT |
| BL413-021-P366 | TTGGAGGTTGATAAATTAAACCTCAGCAGCTTT |
| BL413-024-P378 | CATGATCGATATTAATTGCGTTGCGCTC |
| BL413-024-P379 | GGTCGGCGATGCCGCATAGTTAAGC |
| BL413-024-P380 | CGGCATCGCCGACCAACCGCAAGCG |
| BL413-024-P381 | CAAAGGAAAATTCCAGCTTCCCTGAAACCTTGGACTCC |
| BL413-024-P382 | GCTGGAATTTTCCTTTGCCCTCGGACGAGTGCTG |
| BL413-024-P383 | GCAATTAATATCGATCATGAGCGGAGAATTAAGGGAGTCACG |
| BL413-024-P386 | AAAGAAGAAGAGGAAAGTCGGTGGTAGCGAGCTGATTAAGGAGAAC |
| BL413-024-P387 | ACTTTCCTCTTCTTCTTTGGCTGGAGTGGTCCAGGATTAGATTCAACATC |
| BL413-024-P388 | ATCAATCGGCCGACCAACCGCAAGCG |
| BL413-024-P389 | GTGAGGGCATCGATCATGAGCGGAGAATTAAGGGAGTCAC |
| BL413-024-P390 | TGATCGATGCCCTCACTGGTGAAAAGA |
| BL413-024-P391 | CTTCCTTTCTCGCCACGT |
| BL413-024-P392 | GTGGCGAGAAAGGAAGGG |
| BL413-024-P393 | GGTCGGCCGATTGATGCATGTTGTCAATCAA |
| BL413-024-P429 | AAAGGAATGAAATTCCAGCTTCCCTGAAACCTTG |
| BL413-024-P430 | CTGGAATTTCATTCCTTTGCCCTCGGACG |
| BL413-024-P461 | TCCCACTATCCTTCGCAAGACCCTTCCTCT |
| BL413-024-P462 | AAAAGTCTCAAGTTGCGCAGCCTGAATGG |
| BL413-024-P481 | ATATGATGCTCCAGCCAAAGAAGAAGAGG |
| BL413-024-P482 | CATCACCGACGAGCAAGG |
| BL413-024-P483 | TTGCTCGTCGGTGATGTAC |
| BL413-024-P484 | GGCTGGAGCATCATATTAAGCCTCAGCGATCC |
| BL413-024-P485 | GGGCACAAGTGATAAATTAAACCTCAGCAGCTTTCG |
| BL413-024-P486 | AATTTATCACTTGTGCCCCAGTTTGCT |
| BL413-026-P409 | GCGCAACTTGAGACTTTTCAACAAAGGGTAATATCCG |
| BL413-026-P410 | GGAAAGCCGATAGTGGGATTGTGCGTCATC |
| BL413-027-P603 | TTCTGAGCCTCTTTCCTTCTAATC |
| BL413-027-P604 | CTGTTAATCAGAAAAACTCAGATTAATCGACAAATTCG |
| BL413-027-P605 | GAGTTTTTCTGATTAACAGATGATGTTACAACCAAAGAAGAAAAGGA |
| BL413-027-P606 | TTCAAACCCGGCAGCTTA |
| BL413-027-P607 | AGCTGCCGGGTTTGAAAC |
| BL413-027-P608 | AGGAAAGAGGCTCAGAACCCTTCCCAAATCAGCTTG |
| BL413-037-P681 | ATTGAACTTGACGAACGTTGTCG |
| BL413-037-P697 | CACCCAGCTTTCTTGTACAAAG |
| BL413-037-P698 | CTGTTCTGCTTTTTTGTACAAAGTTGGCA |
| BL413-037-P701 | GTACAAAAAAGCAGAACAGATGGCTGACGCTGTTGTTA |
| BL413-037-P702 | TACAAGAAAGCTGGGTGCTACTGTGTGGCCTTGGATCC |
| BL413-037-P705 | AAACGTCCGCAATGTGTTATT |
| BL413-037-P706 | TGACAGGATATATTGGCGGG |
| BL413-037-P707 | CGCCAATATATCCTGTCAAACACT |
| BL413-037-P708 | AACACATTGCGGACGTTTTTAATG |
