## Supplementary material for "Highlighter: an optogenetic actuator for light-mediated, high resolution gene expression control in plants": S3 Table

**S3 Table**: Synthesized genes used as PCR templates for vector assemblies.

Gene: nlsCcaS(A92V del1_87) codon-optimized for *Arabidopsis*

Legend: Start codons NLS CcaS(A92V del1_87)

Sequence:ATGATGTTACAACCAAAGAAGAAAAGGAAGGTGGGTGGAAGAAGAATAGAAATAAGTATCAAGCAGCAGACACAACGTGAGAGGTTTATCAACCAAATCACACAGCATATCAGACAATCTCTTAATTTGGAGACTGTTTTGAACACTACAGTTGCTGAAGTTAAGACACTTTTGCAGGTTGATAGAGTTCTTATCTATAGAATCTGGCAAGATGGTACAGGATCTGCTATCACTGAGTCTGTTAATGCTAACTACCCTTCTATTTTGGGTAGAACTTTTTCTGATGAGGTTTTCCCAGTTGAATATCATCAAGCTTACACAAAGGGAAAAGTTAGAGCTATTAATGATATCGATCAGGATGATATCGAAATCTGTCTTGCTGATTTCGTTAAACAATTCGGTGTTAAGTCTAAACTTGTTGTTCCTATCTTGCAGCATAATAGAGCTTCTTCTTTGGATAACGAATCTGAGTTTCCATATCTTTGGGGACTTTTGATTACACATCAGTGTGCTTTCACTAGACCTTGGCAACCTTGGGAAGTTGAGCTTATGAAGCAGTTGGCTAACCAAGTTGCTATTGCTATCCAACAGTCTGAGTTGTACGAACAACTTCAACAGTTGAATAAGGATCTTGAGAACAGAGTTGAAAAAAGAACACAACAGTTGGCTGCTACTAATCAGTCTCTTAGGATGGAAATCTCTGAAAGACAAAAGACTGAGGCTGCTTTGAGACATACTAACCATACACTTCAGTCTTTGATTGCTGCTTCTCCTAGAGGTATCTTTACTCTTAATTTGGCTGATCAAATTCAGATCTGGAACCCAACAGCTGAGCGAATCTTCGGATGGACTGAAACAGAGATTATCGCTCATCCTGAGCTTTTGACATCTAACATCCTTTTGGAAGATTACCAACAGTTTAAGCAAAAGGTTCTTTCTGGTATGGTTTCTCCATCTCTTGAGTTGAAGTGTCAGAAGAAAGATGGATCTTGGATTGAAATCGTTTTGTCTGCTGCTCCTCTTTTGGATTCTGAAGAGAACATTGCTGGTCTTGTTGCTGTTGTTGCTGATATCACTGAGCAAAAAAGACAGGCTGAACAAATCAGACTTTTGCAATCTGTTGTTGTTAACACAAACGATGCTGTTGTTATTACTGAAGCTGAACCAATCGATGATCCTGGACCAAGAATCCTTTATGTTAATGAGGCTTTCACTAAGATCACAGGATACACTGCTGAAGAGATGTTGGGAAAGACTCCTAGAGTTCTTCAAGGACCAAAAACTTCAAGAACTGAGTTGGATAGAGTTAGACAGGCTATCTCTCAATGGCAGTCTGTTACAGTTGAAGTTATTAATTACAGAAAGGATGGTTCTGAGTTTTGGGTTGAATTTTCTCTTGTTCCTGTTGCTAACAAAACAGGATTTTACACTCATTGGATTGCTGTTCAAAGAGATGTTACAGAGAGAAGAAGAACTGAAGAGGTTAGACTTGCTTTGGAAAGAGAGAAGGAACTTTCAAGATTGAAGACTAGATTTTTCTCTATGGCTTCTCATGAGTTTAGAACACCACTTTCTACTGCTTTGGCTGCTGCTCAACTTCTTGAAAATTCTGAAGTTGCTTGGCTTGATCCTGATAAGAGATCAAGAAACCTTCATAGAATCCAAAATTCTGTTAAAAACATGGTTCAACTTTTGGATGATATCTTGATTATCAACAGAGCTGAGGCTGGAAAGCTTGAGTTTAATCCAAACTGGCTTGATTTGAAGCTTTTGTTCCAACAGTTCATTGAAGAGATCCAGCTTTCTGTTTCTGATCAATACTACTTCGATTTCATCTGTTCTGCTCAAGATACTAAGGCTCTTGTTGATGAAAGATTGGTTAGATCTATCCTTTCTAATCTTTTGTCTAACGCTATCAAGTACTCTCCTGGAGGTGGACAGATTAAAATCGCTCTTTCTTTGGATTCTGAGCAGATTATCTTCGAAGTTACAGATCAAGGTATTGGAATCTCTCCTGAGGATCAAAAGCAGATCTTTGAACCATTCCATAGAGGAAAGAATGTTAGAAACATTACTGGTACAGGACTTGGTTTGATGGTTGCTAAGAAATGTGTTGATCTTCATTCTGGATCTATCCTTTTGAAGTCTGCTGTGGATCAAGGAACAACTGTGACCATCTGTCTCAAAAGGTACAACCATCTCCCAAGGGCT

Gene: nlsCcaR Codon-optimized for *Arabidopsis*

Legend: Start codons NLS CcaR(del1_3) VP16

Sequence:ATGATGTTGCAGCCTAAAAAGAAGAGAAAAGTTGGTGGTAGAATACTCCTCGTGGAAGATGATTTGCCATTAGCAGAAACCCTCGCAGAAGCTTTGTCTGATCAACTTTACACTGTTGATATTGCTACAGATGCTTCTTTGGCTTGGGATTATGCTTCTAGACTTGAATACGATTTGGTTATTCTTGATGTTATGTTGCCTGAGCTTGATGGAATTACTCTTTGTCAGAAGTGGAGATCTCATTCTTATTTGATGCCAATCCTTATGATGACTGCTAGAGATACAATTAATGATAAGATCACAGGACTTGATGCTGGTGCTGATGATTACGTTGTTAAACCTGTTGATTTGGGTGAACTTTTTGCTAGAGTTAGAGCTCTTTTGAGAAGAGGATGTGCTACTTGTCAACCAGTTTTGGAGTGGGGTCCTATTAGACTTGATCCATCTACTTATGAAGTTTCTTACGATAATGAGGTTTTGTCTCTTACAAGAAAGGAATACTCTATCTTGGAGCTTTTGCTTAGAAACGGAAGAAGAGTTCTTTCTAGATCTATGATCATCGATTCTATCTGGAAGTTGGAGTCTCCTCCAGAAGAGGATACAGTTAAAGTTCATGTTAGATCTTTGAGACAAAAGCTTAAGTCTGCTGGACTTTCTGCTGATGCTATTGAAACTGTTCATGGAATCGGTTACAGATTGGCTAATCTTACAGAGAAGTCTTTGTGTCAGGGAAAGAATTCTTCTGCTCCTCCAACTGATGTTTCTCTTGGAGATGAATTGCATCTTGATGGAGAGGATGTTGCTATGGCTCATGCTGATGCTTTGGATGATTTTGATTTGGATATGCTTGGAGATGGAGATTCTCCTGGACCAGGTTTCACACCTCATGATTCTGCTCCATACGGTGCTCTTGATATGGCTGATTTTGAGTTTGAGCAGATGTTCACTGATGCACTTGGTATTGATGAGTATGGAGGA
