## Supplementary material for "Highlighter: an optogenetic actuator for light-mediated, high resolution gene expression control in plants": S1 Fig

**S1 Fig. Alignment of GAF domains from cyanobacteriochromes, plant phytochromes and bacterial phytochromes.** GAF domains from selected cyanobacteriochromes, plant phytochromes and bacterial phytochromes were aligned to identify functionally conserved amino acid residues involved in chromophore accommodation. In this alignment, the native chromophore of each GAF domain is indicated in parenthesis. The alignment below specifically shows the part of the alignment from the conserved DRV motif to the conserved LWG motif. Residues in the CcaS sequence from Synechocystis sp. PCC6803 (SyCcaS), marked in green, were individually mutated to corresponding amino acids marked in yellow to identify modifications that could improve CcaS photoswitching with PΦB. Details for the cyanobacteriochromes, plant phytochromes and bacterial phytochromes included in the alignment are presented in the table below the alignment. The alignment was performed in CLC Main Workbench 7.6.4 with the following settings: Gap open cost = 10.0, Gap extension cost = 1.0, End gap cost = As any other, Alignment mode = Very accurate, Redo alignments = No, Use fixpoints = No. BV = Biliverdin, PCB = Phycocyanobilin, PVB = Phycoviolobilin and PΦB = Phytochromobilin.

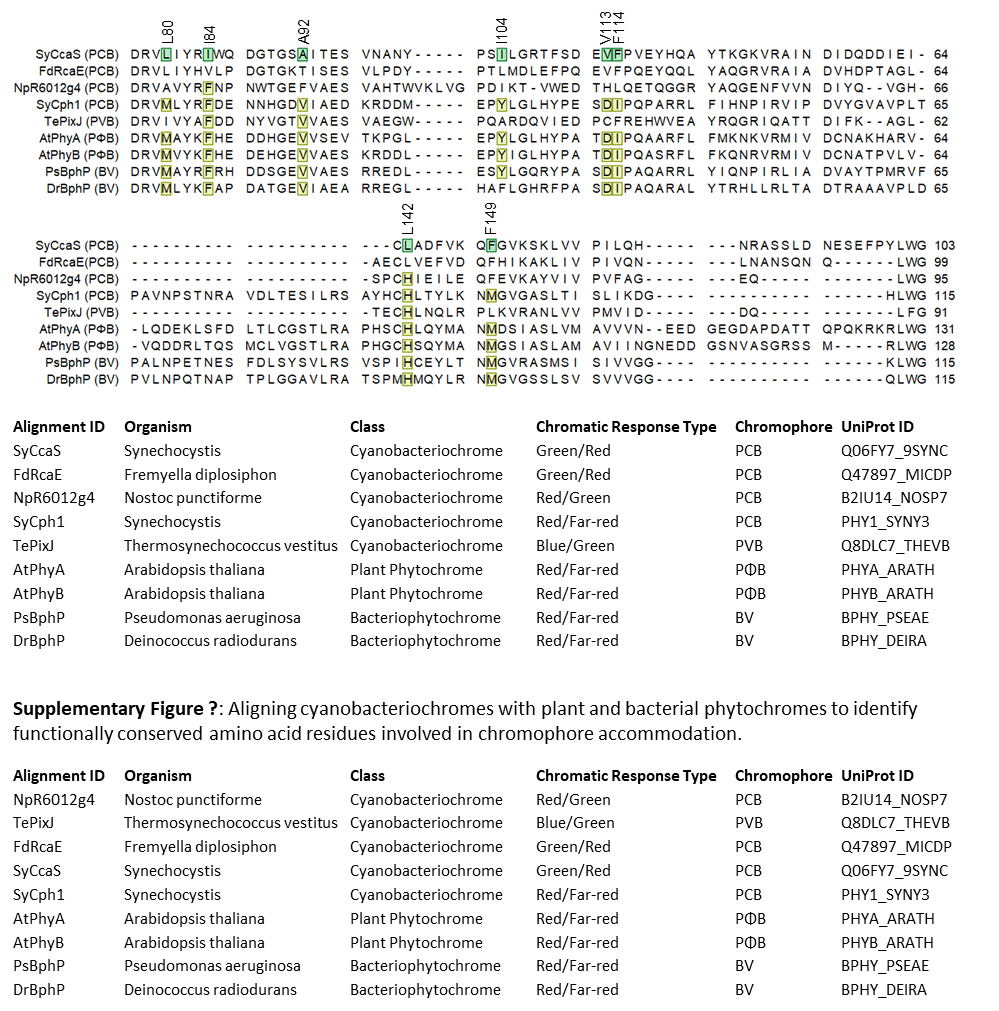
