## Supplementary material for "Highlighter: an optogenetic actuator for light-mediated, high resolution gene expression control in plants": S2 Fig

**S2 Fig. Spectroscopic characterization of holo-CcaS_HL_ purified from PΦB-producing *E. coli*.** (A) Overlay of CcaS_HL_ absorption spectrum for the non-illuminated post-purification sample (black line) and the absorption spectra post 525 nm (red line, red-absorbing state) and 655 nm illumination (green line, green-absorbing state). (B) Difference in absorption spectra between green and red light illuminated holo-CcaS_HL_ isolated from PΦB-producing *E. coli*. (C) Effect of 525 nm illumination on the green-absorbing state of CcaS_HL_. (D) Effect of 655 nm illumination on the red-absorbing state of CcaS_HL_. (E) Effect of 447 nm illumination on the green-absorbing state of CcaS_HL_. (F) Effect of 447 nm illumination on the red-absorbing state of CcaS_HL_; no photoconversion is evident. Absorption spectra in panel (E) were generated with purified holo-CcaS_HL_ from a repeated purification.

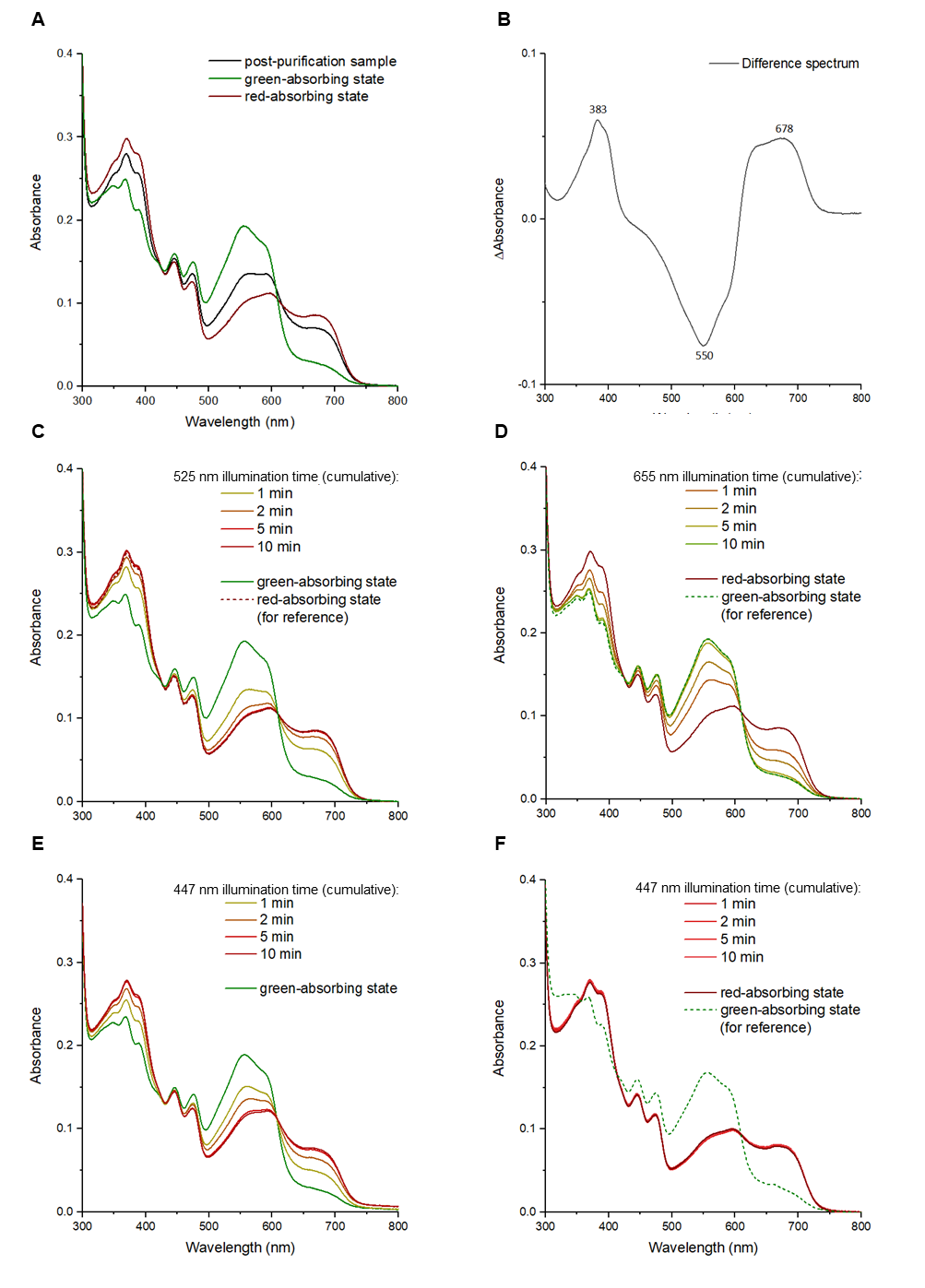
