## Supplementary material for "Highlighter: an optogenetic actuator for light-mediated, high resolution gene expression control in plants": S3 Fig

**S3 Fig. Fluorescence emission spectrum for holo-CcaS_HL_ purified from PΦB-producing *E. coli*.** The fluorescence emission spectrum was acquired between 460nm and 800 nm by exciting the red-absorbing state of holo-CcaS_HL_ purified from PΦB-producing *E. coli* with 445 nm light. Data were acquired from the red absorbing state because blue light illumination causes no further photoisomerization of the PΦB chromophore (S2f Fig). The broad peaks marked at 495 nm and 515 nm are indicative of flavin being bound to the PAS domain of the purified holo-CcaS_HL_, with a striking resemblance to the emission signal from FMN bound to the LOV2 domain of phototropin (Kennis, 2003). The peaks at 628 nm and 666 nm are likely to be from the green-absorbing state of the PΦB chromophore bound to the GAF domain of holo-CcaS_HL_ (Gärtner, 2010).

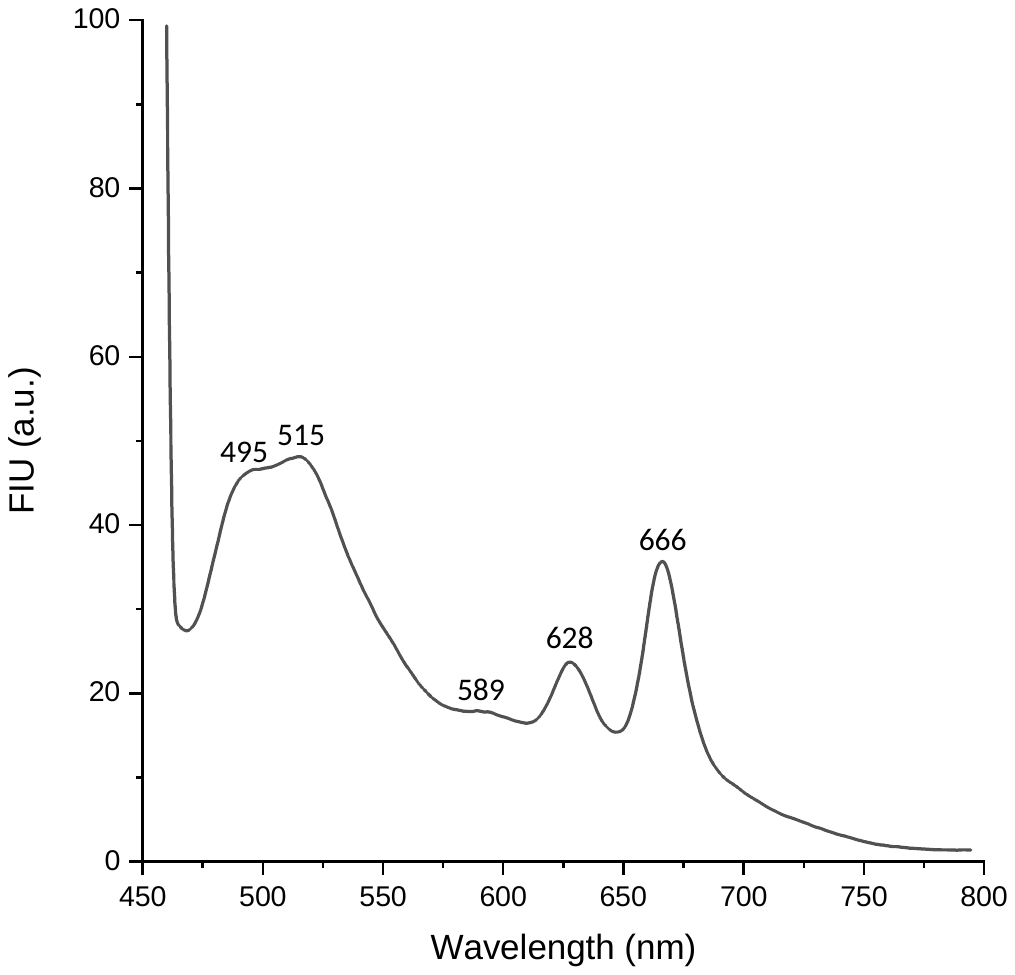
