## Supplementary material for "Highlighter: an optogenetic actuator for light-mediated, high resolution gene expression control in plants": S4 Fig

**S4 Fig**: **Cellular resolution measurements of nuclear YFP/RFP ratios, from Fig 4C, generated by deploying Highlighter(YFP) in transiently transformed *N. benthamiana*.** As describe in Fig 4, a *N. benthamiana* leaf was transiently transformed with Highlighter(YFP) for optogenetic control of YFP expression. The leaf was kept in the dark overnight prior to continuously blue light treatment with LEDs (100 µmol m^-2^ s^-1^) until 2.5 days post infiltration. This was done to keep YFP expression to a minimum prior to light treatment with lasers. Neighboring regions on the transiently transformed leaf were then subjected to blue and red light treatments with lasers. Initiation of laser treatment is defined as 0 h. 442 nm blue and 633 nm red lasers were used to irradiate the area outlined in blue and red in Fig 4A, respectively. The sample received five 7 h light treatments interrupted by confocal imaging. The triangles in the figure below represent individual nuclear YFP/RFP ratios. Their means are graphed in Fig 4C. Blue and red triangles corresponds to nuclei from regions treated with 442 nm blue or 633 nm red lasers, respectively. The blue and red lines trend lines follow the mean values of the blue and red treated nuclei at each time point.

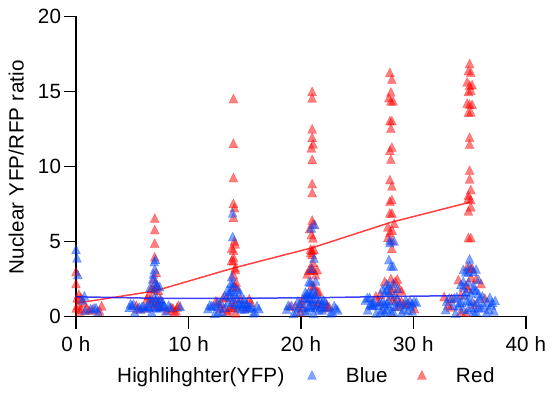
