## Supplementary material for "Highlighter: an optogenetic actuator for light-mediated, high resolution gene expression control in plants": S5 Fig

**S5 Fig**: **Light spectra of LED arrays used for light treating *E. coli* cultures expressing the CcaS&CcaR system variants**. Spectra for the LEDs were recorded with an UPRtek MK350S LED meter. (A) UV. (B) Blue. (C) Green. (D) Yellow. (E) Yellow/Orange. (F) Orange. (G) Red. (H) Far-red.

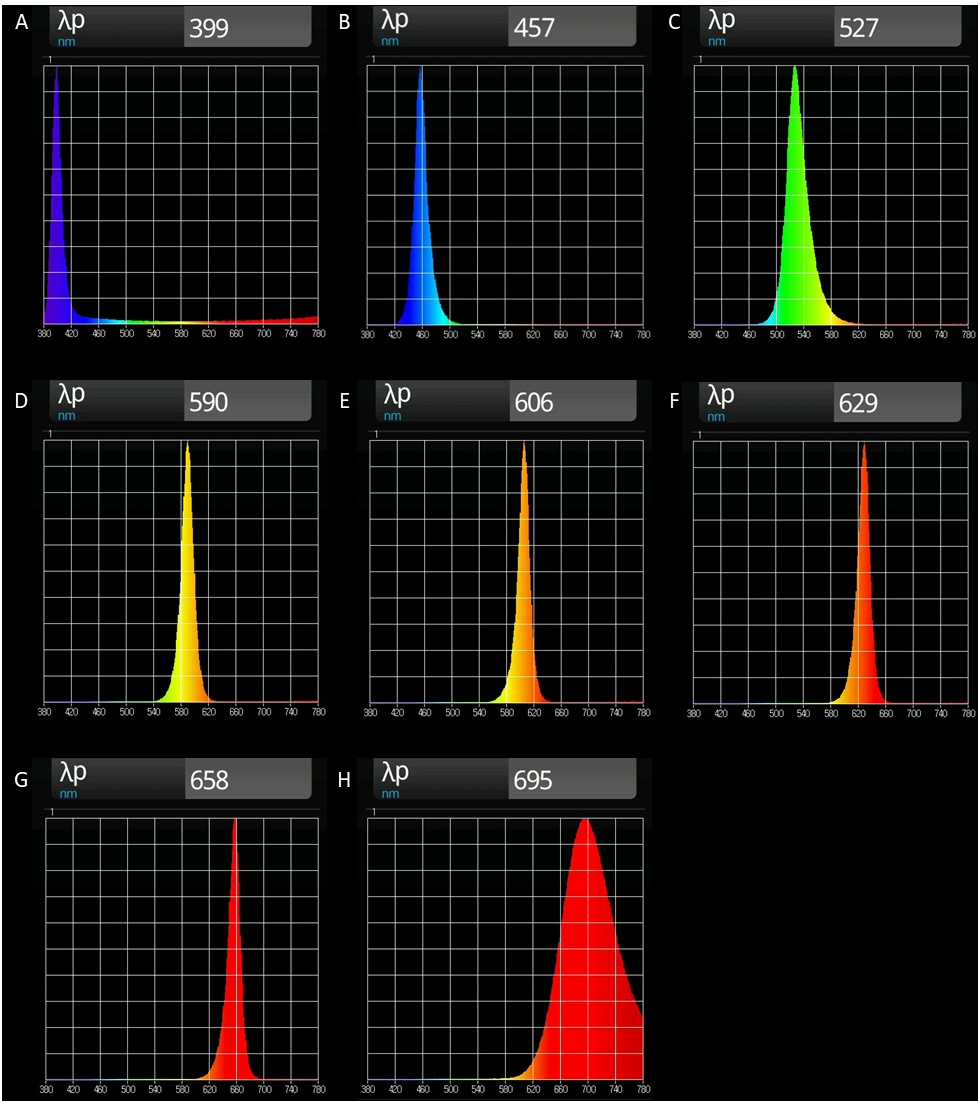

| Color | λp | LED Product |
| --- | --- | --- |
| UV/Blue | 399 nm | UMTMedia 5mm Ultra Bright LEDs (UV) (Amazon) |
| Blue | 457 nm | UMTMedia 5mm Ultra Bright LEDs (Blue) (Amazon) |
| Green | 527 nm | UMTMedia 5mm Ultra Bright LEDs (Green) (Amazon) |
| Yellow | 590 nm | Eddy’s Electronics 5mm Ultra Bright LEDs (Yellow) from (Amazon) |
| Yellow/Orange | 606 nm | UMTMedia 5mm Ultra Bright LEDs (Orange) (Amazon) |
| Orange | 629 nm | UMTMedia 5mm Ultra Bright LEDs (Red) (Amazon) |
| Red | 658 nm | Kingbright 5mm Super Bright LEDs (Red) L-1513SRC-J4 (Rapid Electronics) |
| Far-red | 695 nm | And optoelectronics 3mm LED (Red) AND123R (Farnell) |
