## Supplementary figures and images for "Highlighter: an optogenetic actuator for light-mediated, high resolution gene expression control in plants"

### S6 Fig

**S6 Fig**: **Light spectra of LEDs used for the spectroscopic characterization of holo-CcaS_HL_ in S2 Fig.**

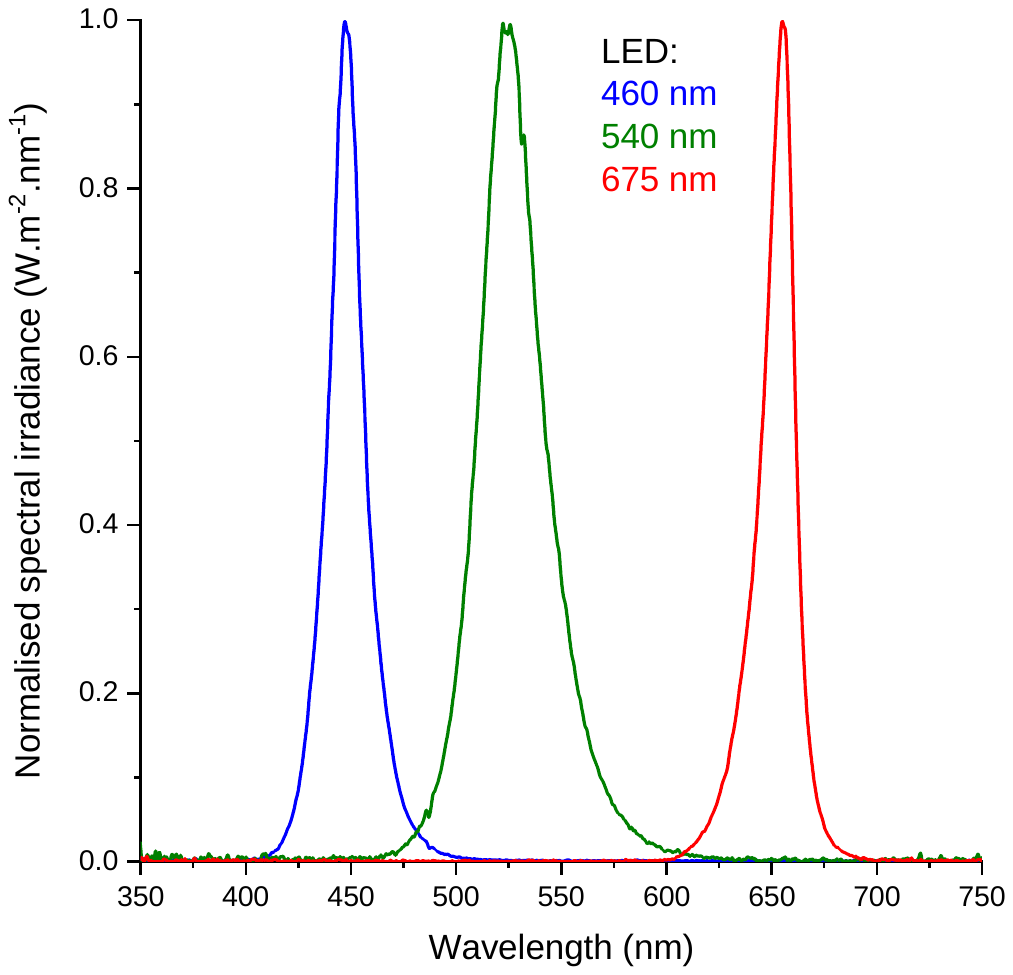
