## Supplementary material for "Highlighter: an optogenetic actuator for light-mediated, high resolution gene expression control in plants": S7 Fig

**S7 Fig**: **Light spectra for Heliospectra RX30 lamps**. Light spectra for light regimes generated with Heliospectra RX30 lamps (Company information in M&M) for light treating transiently transformed / spot infiltrated *N. benthamiana* leaves. Light spectra were recorded with an UPRtek MK350S LED meter. **a**. 450 nm LED channel, **b**. 530 nm LED channel, **c**. 620 nm LED channel, **d**. 660 nm LED channel **e**. 5700 K LED channel (white light LED channel), **f**. Blue enriched white light; 1:1 ration of 450 nm LED channel and 5700 K LED channel, **g**. Green enriched white light; 1:1 ration of 530 nm LED channel and 5700 K LED channel **h**. Orange enriched white light; 1:1 ration of 660 nm LED channel and 5700 K LED channel.

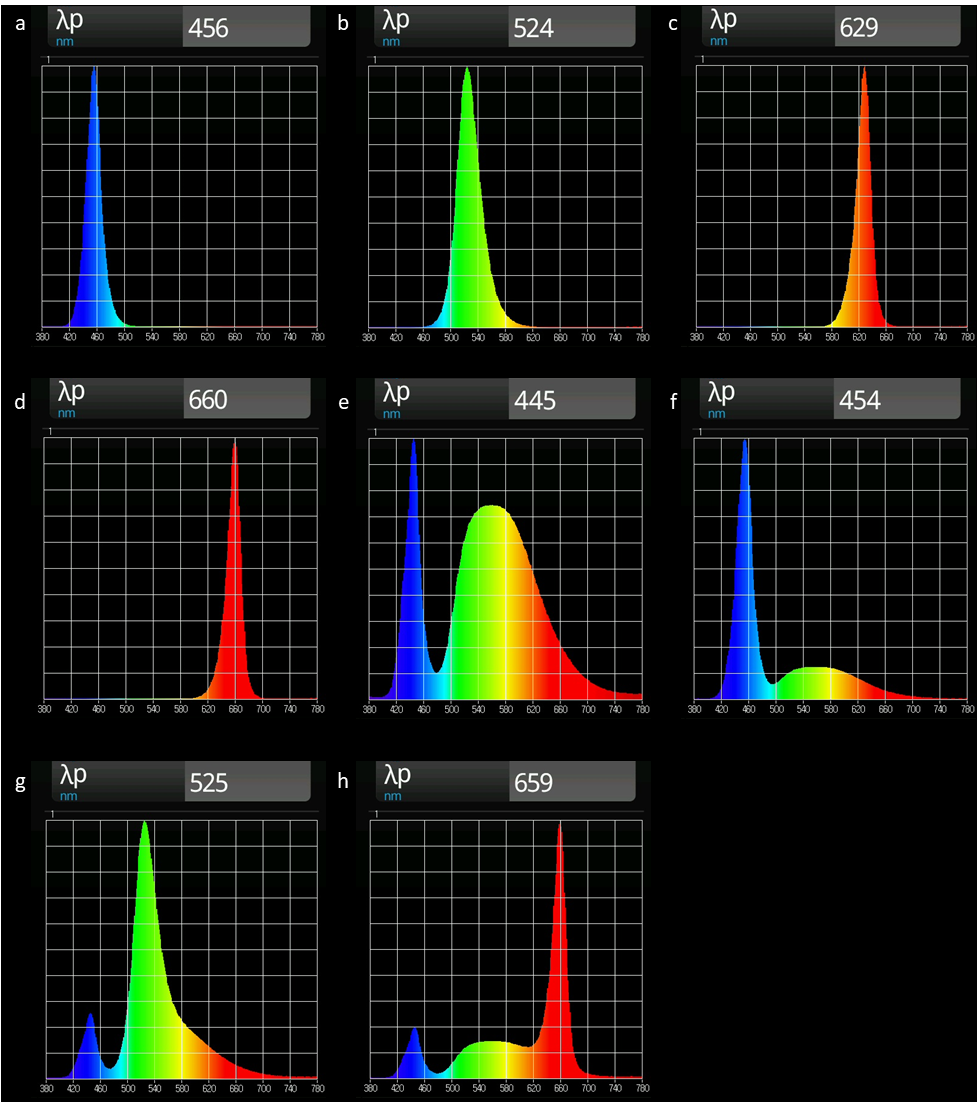
